# Plate-based ISD-SPE enables dual proteome-secretome concentration-response profiling of TLR signalling in iPSC-derived macrophages

**DOI:** 10.64898/2026.07.17.739077

**Authors:** Chloe L. Tayler, Mengchun Li, Charlie Haslam, Kerena Norris, Lee Booty, Rebecca Beveridge, Nicholas J. W. Rattray, Rachel Peltier-Heap

## Abstract

Protein secretion represents a key functional output of cellular signalling, capturing dynamic responses to stimulation and pharmacological perturbation that shape immune behaviour. In macrophages, activation of Toll-like receptors (TLRs) drives tightly regulated secretion programmes that mediate inflammatory responses and provide a biologically meaningful readout of pathway activity. Whilst mass spectrometry (MS)-based secretomics enables unbiased profiling of these processes, broader application in drug discovery remains constrained by sample preparation workflows that limit scalability. Here, we describe a plate-based in-solution digestion and solid-phase extraction (ISD-SPE) workflow that enables 96-well processing of conditioned media for integrated proteome and secretome analysis from the same sample well. Benchmarking against a precipitation-based approach demonstrated comparable proteomic depth with improved quantitative reproducibility and robust performance across multiple plates. Coupled with dia-PASEF acquisition, this workflow enabled in-depth profiling of macrophage responses to TLR activation, resolving receptor-specific secretory programmes following TLR3, TLR4 and TLR7/8 activation. Extension of the approach to concentration-response studies enabled quantitative characterisation of pharmacological perturbation across intracellular and extracellular protein landscapes, revealing both shared and compartment-specific responses to TLR inhibition, as well as differences in apparent potency linked to secretion dynamics. Together, this workflow provides a scalable strategy for integrated analysis of intracellular signalling and downstream protein secretion, enabling systems-level characterisation of inflammatory responses and compound mechanisms of action.

## Introduction

The active secretion of proteins into the extracellular space is a fundamental mechanism by which cells coordinate intercellular communication and regulate inflammatory responses (1, 2). Collectively known as the secretome (3), these proteins encompass a diverse range of functional classes, including cytokines, chemokines and growth factors (4, 5). In the context of innate immunity, macrophages utilise tightly regulated secretion programmes in response to environmental stimuli, including viruses, bacteria and endogenous cellular damage, with activation of pattern recognition receptors (PRRs), including Toll-like receptors (TLRs), driving downstream signalling cascades that result in the release of pro-inflammatory cytokines and interferon-stimulated gene products (ISGs) (6, 7). These secreted proteins represent key functional outputs of immune activation and play central roles in shaping both local and systemic immune responses.

The secretome represents a highly dynamic and pharmacologically relevant subset of the proteome, reflecting both the activation state of a cell and its interactions with the surrounding microenvironment (8, 9). In the context of inflammatory disease and drug discovery, secreted proteins serve as key functional outputs of cellular signalling, acting as mediators of pathology whilst also providing a rich source of biomarkers for monitoring therapeutic response (10, 11). Many clinically validated drug targets, including tumour necrosis factor (TNF) and members of the interleukin family, are themselves secreted proteins (12–14). Profiling changes in the secretome therefore provides a functional, systems-level readout of pathway engagement and cellular responses to perturbation. Consequently, robust technologies for the detection and quantification of secreted proteins are essential for enabling translational research and drug discovery efforts. Immunoassay platforms, including enzyme-linked immunosorbent assays (ELISA) and multiplex bead-based alternatives, remain the gold standard for measuring secreted proteins due to their sensitivity and ease of use (15–17). Technologies such as proximity extension assays (PEA; Olink) and nucleobase-enabled localised immunoassays with spectral addressing (nELISA; Nomic Bio) have further extended these approaches through enhanced multiplexing and improved quantitative performance (18, 19). However, these methods rely on predefined antibody panels and prior target knowledge, limiting their ability to enable unbiased discovery.

Mass spectrometry (MS)-based proteomics offers an untargeted approach for protein identification and quantification, enabling detection of thousands of proteins per sample across a broad dynamic range (20, 21). Technological advances in data-independent acquisition (DIA) strategies, together with streamlined sample preparation workflows and deep learning-based data analysis tools, have substantially improved both throughput and proteomic depth (22–24). In particular, the integration of trapped ion mobility spectrometry (TIMS) with parallel accumulation-serial fragmentation (PASEF) has proven transformative, enabling high-speed acquisition times and near 100% duty cycles without compromising sensitivity (25, 26). We recently demonstrated the application of this platform to secretomics, establishing a robust dia-PASEF workflow that reproducibly quantifies over 900 proteins from iPSC-derived macrophage supernatants in under 15 minutes of acquisition time (27). This approach enabled the resolution of stimulus-specific inflammatory signatures, identification of biological features not captured by targeted immunoassay panels, and characterisation of temporal secretion dynamics during macrophage activation.

Despite these capabilities, a key limitation of conventional secretomics workflows is their reliance on protein precipitation methods, such as acetone precipitation, for sample preparation. Whilst effective for removing media additives and concentrating proteins from dilute supernatants (28), precipitation-based approaches introduce practical challenges for scale. Resulting protein pellets can be difficult to reproducibly resolubilise, recovery from low-input samples is often inconsistent, and the use of volatile organic solvents limits compatibility with automated, plate-based liquid handling systems (28–30). Collectively, these constraints restrict the throughput and scalability of secretomics in early-stage drug discovery, where large-scale, multi-condition studies are routinely employed. There is therefore a clear need for miniaturised, plate-compatible sample preparation strategies that preserve proteomic depth and quantitative reproducibility whilst enabling efficient processing of large sample cohorts.

To address these limitations, we present a plate-based in-solution digestion and solid-phase extraction (ISD-SPE) workflow for secretomics, compatible with standard 96-well formats. We demonstrate that this approach achieves proteomic coverage comparable to conventional precipitation methods whilst improving quantitative reproducibility and scalability. Integrated with our previously described dia-PASEF platform, the workflow is fully compatible with low-input 96-well cell culture systems and enables parallel profiling of the proteome and secretome from the same well, providing complementary intracellular and extracellular readouts within a single experiment.

The utility of this platform was evaluated across two complementary discovery contexts. Time-resolved secretome profiling of macrophage responses to TLR3, TLR4, and TLR7/8 activation resolved receptor-specific patterns of cytokine, chemokine and ISG secretion over an 8-hour period, including distinct interferon-driven responses associated with TIR-domain-containing adapter-inducing interferon-β (TRIF)-dependent signalling. In parallel, a global concentration-response framework was developed to characterise concentration-dependent effects of selective TLR4 and TLR7/8 inhibition across the proteome and secretome. This analysis identified concentration-responsive cytokines and chemokines alongside corresponding intracellular changes, revealing differences in potency and regulatory dynamics between compartments. Collectively, this integrated approach enables resolution of pathway- and compartment-specific responses to pharmacological perturbation that are not fully captured by either dataset alone, highlighting the potential of this platform for scalable, systems-level interrogation of inflammatory signalling in drug discovery.

## Experimental Procedures

### Macrophage differentiation and maintenance

Human induced pluripotent stem cell (iPSC) lines UKBi006-A, 2094BW-A2:7, 6732BW-M3:2 and 7980BW-S2:1 were supplied by the European bank for Induced Pluripotent Stem Cells (EBiSC) and the use of samples received Ethics Committee approval (CEI-117-2357). Cell expansion, embryoid body formation and monocyte-like production were carried out according to the protocol previously described by Armesilla-Diaz *et al* (31). Monocyte precursors were counted (NucleoCounter^®^, NC-200, Via1-Cassette^™^) and 16,000 cells/well were seeded in 96-well culture plates with macrophage differentiation medium, which consisted of RPMI (with GlutaMAX^™^ supplement, Gibco^™^), 10% Fetal Bovine Serum (FBS, Gibco^™^) and 100 ng/mL Macrophage Colony-Stimulating Factor (M-CSF, PeproTech). After 6 days, cells were visually inspected to confirm differentiation into resting (M0) macrophages. All cultures were maintained at 37°C in a humidified atmosphere of 5% CO_2_.

### Cell treatments and preparation of secretome samples

All experiments were conducted in 96-well plates. Washing steps were carried out using Dulbecco’s Phosphate Buffered Saline (DPBS, Sigma-Aldrich), and treatments were applied using media prewarmed to 37°C. Macrophage stimulation reagents were sourced from InvivoGen at the highest available purity. For method validation experiments, differentiation medium was removed and cells were washed twice before the addition of OptiMEM^™^ reduced serum medium (Gibco^™^) supplemented with 100 ng/mL M-CSF and lipopolysaccharide (LPS) at 100 ng/mL. Control samples received the reduced serum medium supplemented with M-CSF, but no additional stimulation. Cells were incubated for 4 hours before the supernatants were carefully harvested and transferred to 96-well LoBind plates (Eppendorf). Supernatants were centrifuged at 1500 *g*, 4°C for 30 minutes to pellet cell debris. Clarified supernatants were then transferred to a fresh plate and stored at −80°C. Remaining adherent cells were washed twice before the addition of lysis buffer (5% (v/v) sodium dodecyl sulphate (SDS), 100 mM triethylammonium acetate buffer (TEAB, Sigma-Aldrich) (aq)) containing 25 U of Benzonase^®^ Nuclease (Sigma-Aldrich). After 30 minutes, lysates were harvested and stored in a 96-well LoBind plate at −80°C.

For interval-based kinetics experiments, differentiation medium was exchanged with fresh differentiation medium supplemented with TLR agonists (polyinosinic-polycytidylic acid (Poly(I:C)), 100 ng/mL; LPS, 100 ng/mL; resiquimod (R848), 1000 ng/mL). Macrophages were incubated for the indicated time points following stimulation. Two hours before the end of each time point, stimulation medium was removed and cells were washed twice. An additional two hour incubation was then performed in OptiMEM^™^ supplemented with 100 ng/mL M-CSF and the corresponding TLR agonist. Supernatants and cell lysates were subsequently harvested, processed as previously described, and stored at −80°C until further analysis.

For concentration-response experiments, small molecule inhibitors resatorvid (TAK-242; TLR4) and MHV370 (TLR7/8) were tested using an 11-point 1:3 dilution series, with a top concentration of 10 µM and matched DMSO vehicle controls included (final DMSO concentration 1%). Cells were pre-treated with inhibitors for 2 hours in macrophage differentiation medium prior to TLR stimulation. Following inhibitor pre-treatment, the appropriate TLR agonist was added (LPS at 100 ng/mL for TLR4, or R848 at 1000 ng/mL for TLR7/8), and cells were incubated for 4 hours in macrophage differentiation medium. Medium was then removed, cells were washed twice, and a second 4-hour incubation was performed in OptiMEM^™^ supplemented with 100 ng/mL M-CSF, the respective agonist, and the appropriate concentration of inhibitor, to enable secretome collection. At the end of each OptiMEM^™^ incubation, cell culture supernatants and cell lysates were harvested as described above and stored at −80°C until further analysis.

### Live-cell imaging assay

Cells were imaged following a 4-hour incubation in Opti-MEM^™^ to assess viability and well-to-well distribution. The supernatant was removed, and cells were washed twice before incubation with Hoechst 33342 (1:1000 dilution, Invitrogen) for 45 minutes at 37°C, 5% CO_2_, to stain nuclei. Dye was subsequently removed, and cells then incubated with Calcein AM and BOBO-3 Iodide from the LIVE/DEAD^™^ imaging kit (Invitrogen) according to the manufacturer’s protocol to label viable and non-viable cells, respectively. Imaging analysis was performed using a Yokogawa CV8000 system with a 10X/0.45NA objective lens. Detailed parameters of the SIMA analysis pipeline have been provided in Table S1.

### Acetone precipitation of supernatants

Using 1.5 mL LoBind tubes (Eppendorf), 400 µL of ice-cold acetone was added to 100 µL of supernatant and vortexed briefly before incubation at −20°C for 60 minutes. Samples were subsequently centrifuged at 10,000 *g*, 4°C for 30 minutes before aspiration of the supernatant. The resulting protein pellets were allowed to air dry for 30 minutes before resuspension in 50 µL of lysis buffer.

### In-solution digestion and solid-phase extraction (ISD-SPE) of supernatants

Supernatant samples were processed using a plate-based in-solution digestion and solid phase extraction (ISD-SPE) workflow. Frozen 96-well plates containing conditioned media were thawed, and samples were adjusted to 50 mM TEAB. Proteins were reduced with 10 mM tris(2-carboxyethyl)phosphine (TCEP) for 30 minutes at room temperature, followed by alkylation with 10 mM iodoacetamide (IAA) for 30 minutes in the dark at room temperature. Proteins were digested by the addition of 100 ng of Trypsin/LysC and incubated for 18 hours at 37°C. Digestion was quenched by acidification with a final concentration of 0.1% (v/v) trifluoroacetic acid (TFA). Resulting peptides were desalted using a Sep-Pak tC18 96-well µElution plate (Waters), with all conditioning, sample loading, washing and elution steps performed by centrifugation at 200 *g* for 5 minutes. The Sep-Pak plate was first conditioned and equilibrated with 0.1% (v/v) formic acid in acetonitrile (buffer B) and 0.1% (v/v) formic acid in water (buffer A) prior to sample loading. Bound peptides were washed three times with 150 µL of buffer A and eluted twice with 150 µL of 50% (v/v) acetonitrile containing 0.1% formic acid (aq).

### Proteomic digestion of cell lysates and acetone extracted protein secretions

Cell lysates and acetone extracted protein secretions were processed using the S-Trap^™^ digestion protocol published by ProtiFi (32). In brief, proteins were reduced with 10 mM TCEP for 30 minutes at room temperature, before alkylation with 10 mM IAA for 30 minutes in the dark at room temperature. The alkylation reaction was quenched with the addition of phosphoric acid to a final concentration of 12.5% (v/v) and gentle vortexing. Acidified samples were diluted with binding buffer (100 mM TEAB in 90% methanol (aq)), loaded onto an S-Trap^™^ 96-well digest plate (ProtiFi) and centrifuged at 1500 *g* for 2 minutes. Samples were washed three times with binding buffer before the addition of 400 ng of Trypsin/LysC in 125 µL of 50 mM TEAB (aq) to each well. Samples were incubated at 37°C for 18 hours before elution from the digest plate. Resulting peptides were dried under vacuum centrifugation and stored at −80°C ready for LC-MS/MS analysis.

### Sample preparation for mass spectrometry

Lyophilised supernatant and cell lysate samples were resuspended in 100 µL of 0.1% (v/v) formic acid (aq), after which 10% of each sample was subsequently loaded onto Evotips (Evosep Biosystems) according to the manufacturer’s instructions for LC-MS/MS analysis.

### LC-MS/MS analysis

All samples were analysed using an Evosep One (Evosep Biosystems) LC system coupled with a TimsTOF HT mass spectrometer (Bruker Daltonics). Cell lysate and supernatant samples were separated using the 60 or 100 samples per day (SPD) methods, respectively. Both gradient methods were run using the recommended Evosep Performance column (C18, 1.5 µm, 8cm x 150µm, EV1109), maintained at 50°C. The analytical column was connected to a nano electrospray ion source (CaptiveSpray 2 source; Bruker Daltonics) with a 10 µm ID fused silica emitter (Bruker Daltonics). The mobile phases consisted of 0.1% (v/v) formic acid (aq) as buffer A and 0.1% (v/v) formic acid in acetonitrile as buffer B. All solvents were of LC-MS grade and sourced from Fisher Scientific.

The mass spectrometer was operated in dia-PASEF mode for all measurements, with an ion mobility range from 1/K0 = 1.6 to 0.6 Vs cm^-2^. Ion accumulation and ramp times were set to 100 ms each. Mass spectra were acquired from 100 – 1700 *m/z*. Eight dia-PASEF scans per TIMS-MS scan were used to give a duty cycle of 0.96 seconds. The dia-PASEF method was optimised using py_diAID (33) to cover a range of 300 – 1200 *m/z*, including 3 ion mobility windows per dia-PASEF scan with variable isolation widths adjusted to the precursor densities (Table S2).

### Proteomics data analysis

All raw MS data files were analysed using Data Independent Acquisition by Neural Networks (DIA-NN) Enterprise v2.2.0 (24), using default precursor ion generation settings and an in silico predicted spectral library allowing for cysteine carbamidomethylation, N-terminal methionine excision and 1 missed cleavage. The spectral library was generated from a human reference database (UniProt 2023 release, 20,399 sequences). Algorithm settings were as follows: Scoring, ‘Generic’, Proteotypicity, ‘Protein names (from FASTA)’, Machine learning, ‘NNs (cross-validated)’, Quantification strategy, ‘QuantUMS (high precision)’, Cross-run normalisation, ‘RT-dependent’, Library generation, ‘IDs, RT & IM profiling’, Speed and RAM usage, ‘Optimal results’. Both mass accuracy and MS1 accuracy were set to 0 for automatic inference. ‘Match between runs’ and ‘Protein inference’ were enabled. The precursor and protein-level false discovery rates (FDR) were set to 1%.

### Proteomics statistical analysis

Statistical analyses were performed using the DIA-NN protein group matrix report files. Sample load normalisation was first performed on the raw data using a variance stabilising normalisation (VSN), assuming that there is a proportional relationship between the total protein from any sample and the total raw signal associated with that sample (34). Whole cell lysate samples were also batch corrected to remove individual donor effects using the removeBatchEffect function of the Limma R package (v3.66.0) (35).

Log_2_-transformed, normalised protein intensities were used for differential expression analysis in Perseus (v1.6.15.0). Data were first filtered for contaminants and 66% valid values in at least one experimental condition. Missing values were imputed drawing from a normal distribution (width 0.3 and downshift 1.8). Two-sample t-tests were applied between groups to identify significantly regulated proteins, with a p-value cutoff of 0.05.

Concentration-response curves were fitted using imputed log_2_-transformed intensity values. A four-parameter log-logistic model was fitted using the drc R package (v3.0.1) (36), allowing all parameters to be freely estimated. A null model assuming no concentration dependence was fitted for comparison. Model fit was evaluated by comparing Akaike’s Information Criterion (AIC) (37), calculated using the stats R package (v4.5.1). In addition, an R^2^ value was computed to quantify the improvement of the log-logistic model over the flat model, defined as:

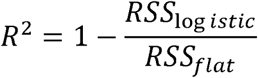

Proteins with an R^2^≥ 0.5, effect size > 0.5 and effective concentration at half maximum (EC_50_) within the experimental concentration range were reported as concentration responsive.

Gene ontology (GO) and pathway enrichment analyses were performed using the Database for Annotation, Visualisation and Integrated Discovery (DAVID) and STRING (38, 39). The screening criterion for a significantly regulated ontology or pathway was a p-value of ≤ 0.05.

### Data visualisation

Biorender (https://www.biorender.com/), Cytoscape v3.10.4 (https://cytoscape.org/), GraphPad Prism v9.3.1 (https://www.graphpad.com/) and R v4.5.1 (https://www.r-project.org/) were used for workflow and data illustrations.

### Experimental design and statistical rationale

The mass spectrometry proteomics and secretomics data have been deposited in the ProteomeXchange Consortium via the PRIDE partner repository with the dataset identifier PXD081205. All experiments were performed using iPSC-derived macrophages.

Benchmarking of sample preparation workflows was performed using 48 replicates per workflow (24 resting, M0; 24 pro-inflammatory, M1) for both acetone precipitation and ISD-SPE, enabling direct comparison of technical variability alongside retention of biological signatures. ISD-SPE robustness and scalability were further assessed by quantifying intra-plate variability across a full 96-well plate with a balanced M0/M1 layout, and inter-plate variability using the same design across two plates. Time-resolved kinetic experiments were conducted using three biological replicates per condition, which were sufficient to capture consistent temporal secretion. Concentration-response experiments comprised three biological and three analytical replicates per condition, enabling robust estimation of biological and technical variability and validation of the data processing pipeline. Analytical replicates were averaged per donor for downstream analysis. All MS run orders were randomised to avoid potential carryover effects or any similar biases.

## Results

### Benchmarking and analytical validation of a 96-well ISD-SPE workflow for secretomics

We first sought to translate our recently established dia-PASEF secretomics workflow into a miniaturised, plate-based format suitable for scalable applications. In our previous work, we demonstrated that this platform enables rapid, unbiased profiling of the human secretome, reproducibly quantifying over 900 proteins in an iPSC-derived macrophage model. Whilst showing good overall agreement with a conventional antibody-based panel, our approach additionally captured stimulus-specific biology that was not detected by these targeted assays. Furthermore, we used this platform to resolve dynamic secretion trajectories, enabling the distinction between acute and chronic inflammatory states (27). Building on these observations, we employed the same iPSC-derived macrophage model from three human donors and stimulated with lipopolysaccharide (LPS) for four hours to induce a pro-inflammatory (M1) phenotype (Figure 1A). Cells were cultured for 6 days in a 96-well format, with brightfield microscopy confirming morphological changes consistent with the transition from monocyte precursor to the resting macrophage (M0) phenotype, as described in our previous work.

**Figure 1.**
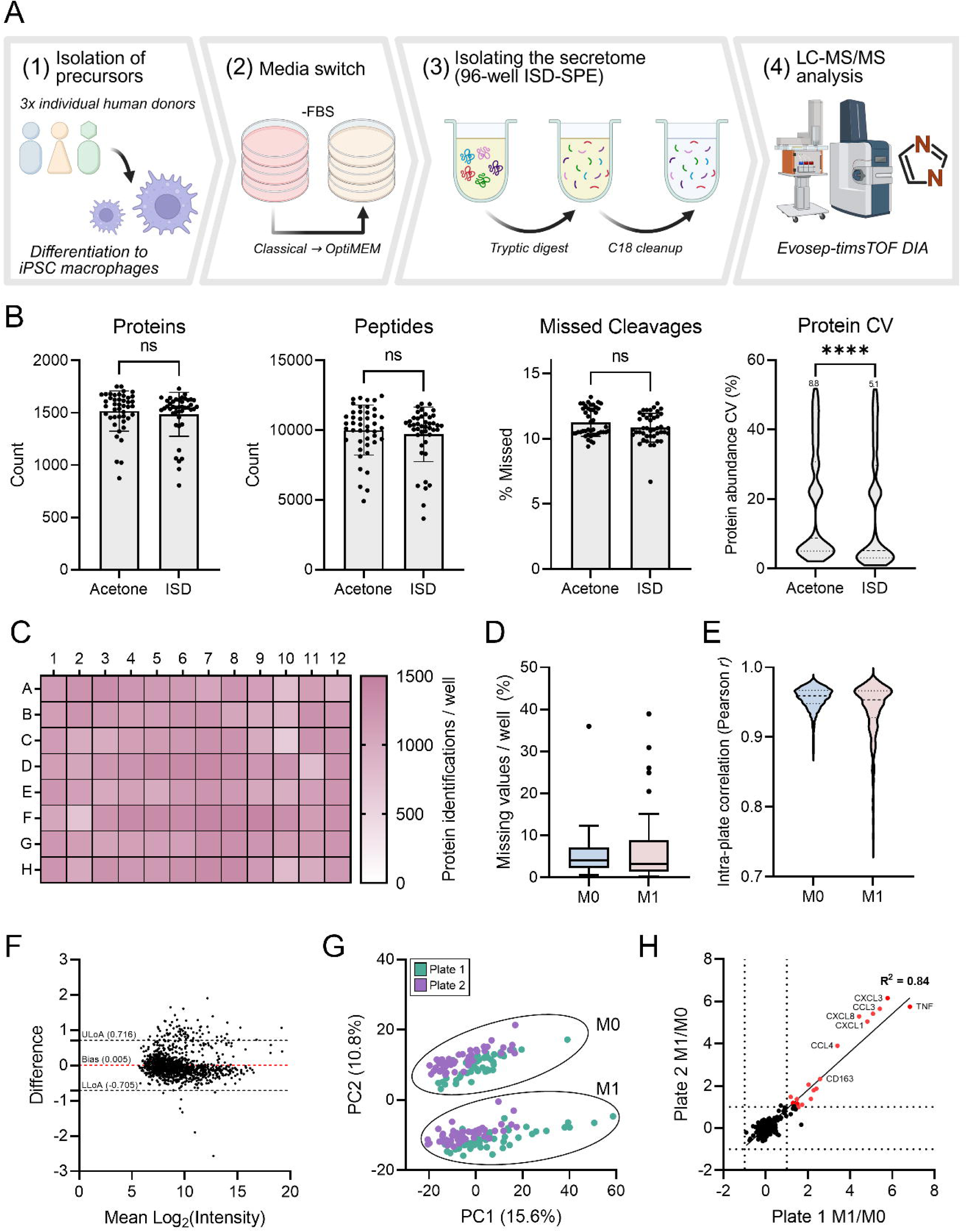
ISD-SPE enables robust and reproducible secretome analysis in a 96-well format. (A) Schematic of the MS-based ISD-SPE workflow, including sample preparation, tryptic digestion, C18 solid-phase extraction, and dia-PASEF acquisition. (B) Comparison between acetone precipitation and ISD-SPE. (C – E) Intra-plate reproducibility. (C) Heatmap of protein identifications demonstrating uniform proteome coverage across the plate (D) Distribution of missing values per well show low data incompleteness across M0 and M1 conditions. (E) Pairwise Pearson correlations within each condition indicate high reproducibility (median r ∼ 0.95). (F – H) Inter-plate reproducibility. (F) Bland-Altman plot comparing protein abundances between plates. Differences are centred around zero (bias = 0.005), with 95% of proteins within the limits of agreement, indicating minimal systematic bias. (G) Principal component analysis (PCA) of secretomes from two independent plates shows clustering by biological condition (M0 and M1). Samples are coloured by plate to illustrate minimal contribution of batch-driven variation to the overall secretome. (H) Log_2_(fold change) comparison between plates; fold changes calculated relative to M0 controls and filtered for p < 0.05.

To minimise background interference from highly abundant serum proteins, cells were switched to Opti-MEM™ reduced serum medium at the point of stimulation. Cultures were imaged following supernatant harvest to assess well-to-well seeding uniformity and cell viability. Fluorescence microscopy using Hoechst, Calcein AM and BOBO-3 iodide staining demonstrated consistent cell distribution across the plate and high viability (≥95%) in both M0 and M1 populations (Figure S1A – B). These results confirm the robustness of the 96-well cell culture system and indicate that neither the reduced serum conditions nor LPS stimulation adversely affected cell health after four hours.

Whilst acetone precipitation is widely used to extract proteins from cell culture supernatants, its application to larger scale studies remains challenging as resulting protein pellets can be difficult to resolubilise, recovery from dilute samples is often inconsistent, and the handling of volatile solvents is not readily compatible with automated workflows (30). To address these limitations, we compared conventional acetone precipitation with a plate-based in-solution digestion and solid-phase extraction (ISD-SPE) approach. Following supernatant harvest, proteins were either extracted by acetone precipitation prior to tryptic digestion or processed using the ISD-SPE workflow before subsequent DIA analysis (Figure 1A). Secretomic analysis of the supernatants prepared by each method revealed comparable peptide and protein identifications, alongside similar proportions of missed cleavages and a high degree of overlap in the proteins identified (Figure 1B, Figure S2A). Proteins uniquely detected in each dataset largely carried intracellular annotations and lacked clear biological relevance (Table S3). Despite the overall similarity between the two datasets, protein-level coefficient of variation (%CV) analysis demonstrated improved quantitative reproducibility for the ISD-SPE workflow compared to acetone precipitation, with median CVs of 5.1% and 8.8%, respectively (Figure 1B). Both approaches captured a robust M1-characteristic secretome following LPS stimulation, marked by the significant upregulation of pro-inflammatory cytokines and chemokines. We observed a strong agreement in the fold changes of the significantly upregulated proteins between workflows (R^2^ = 0.81, Figure S2B), indicating that neither approach alters the underlying biology of the samples nor introduces bias in protein recovery. Gene Ontology (GO) Cellular Component (CC) enrichment analysis of the ISD-SPE dataset showed significant enrichment of extracellularly annotated and cytosolic proteins (Figure S2C – D), indicating that this workflow produces supernatants enriched for secreted and secretion-associated proteins with minimal contribution from intracellular contaminants. Based on these results, and the streamlined, miniaturised nature of the ISD-SPE workflow, we proceeded to evaluate its robustness and suitability for larger-scale, multi-plate applications.

We next assessed the intra-plate reproducibility of the ISD-SPE workflow across a full 96-well plate. Protein identifications were consistent across all wells, with a median of 1189 proteins identified and a CV of 13%, and no evidence of edge effects across the plate (Figure 1C). Missing values remained low and comparable between M0 and M1 populations (Figure 1D), with median values of 4.0% and 3.2%, respectively, indicating minimal data incompleteness. To evaluate quantitative reproducibility, pairwise Pearson correlations were calculated between wells within each condition (Figure 1E). Replicates showed excellent agreement, with median correlation coefficients exceeding 0.95 in both groups.

Reproducibility across multiple plates is essential for scalable, high throughput secretomics applications. We therefore proceeded to evaluate the inter-plate performance of the ISD-SPE workflow using two independent 96-well plates. Comparison of protein abundances between plates using a Bland-Altman approach showed differences centred around zero (bias = 0.005), with more than 95% of proteins falling within the limits of agreement, indicating minimal systematic bias (Figure 1F). Proteins outside the limits of agreement were largely extracellular and media-associated, with no clear biological relevance (Table S4). Principal component analysis (PCA) of secretomes from both plates showed distinct clustering by biological phenotype rather than by plate (Figure 1G). Consistent with this outcome, strong correlations were observed between plates (median = 0.94; Figure S3). Furthermore, strong agreement in the fold changes of significantly upregulated proteins confirmed that biologically relevant differences were conserved across experiments (R^2^ = 0.84; Figure 1H).

Collectively, these data demonstrate that our ISD-SPE workflow delivers robust, reproducible and quantitatively consistent performance both within and across plates, supporting its application in larger-scale studies. This workflow was therefore adopted for all subsequent experiments and applied to both time-resolved and concentration-dependent perturbations to interrogate Toll-like receptor (TLR) signalling.

### Time-resolved secretomics reveals receptor-specific responses to TLR activation

Macrophage responses to pathogen-associated and self-derived stimuli are mediated through the activation of distinct Toll-like receptor (TLR) signalling pathways, resulting in the secretion of pro-inflammatory cytokines, chemokines and interferon-stimulated genes (ISGs) (40, 41). Whilst members of the TLR family exhibit a high degree of specificity in ligand recognition, their signalling is largely mediated by two adaptor molecules, myeloid differentiation primary response protein 88 (MyD88) and TIR domain-containing adaptor molecule 1 (TRIF), leading to substantial overlap in downstream responses (Figure 2A) (42, 43). Nevertheless, activation of distinct TLRs has been shown to result in highly overlapping yet differential secretory profiles, reflecting subtle differences in downstream pathway engagement (44). To investigate this further, we applied our optimised sample preparation workflow and dia-PASEF platform to profile the time-resolved secretome of iPSC-derived macrophages following activation of TLR3 (Poly(I:C), 100 ng/mL), TLR4 (LPS, 100 ng/mL), or TLR7/8 (R848, 1 µg/mL) over an 8-hour period, with media exchanged every two hours to capture discrete secretion intervals.

**Figure 2.**
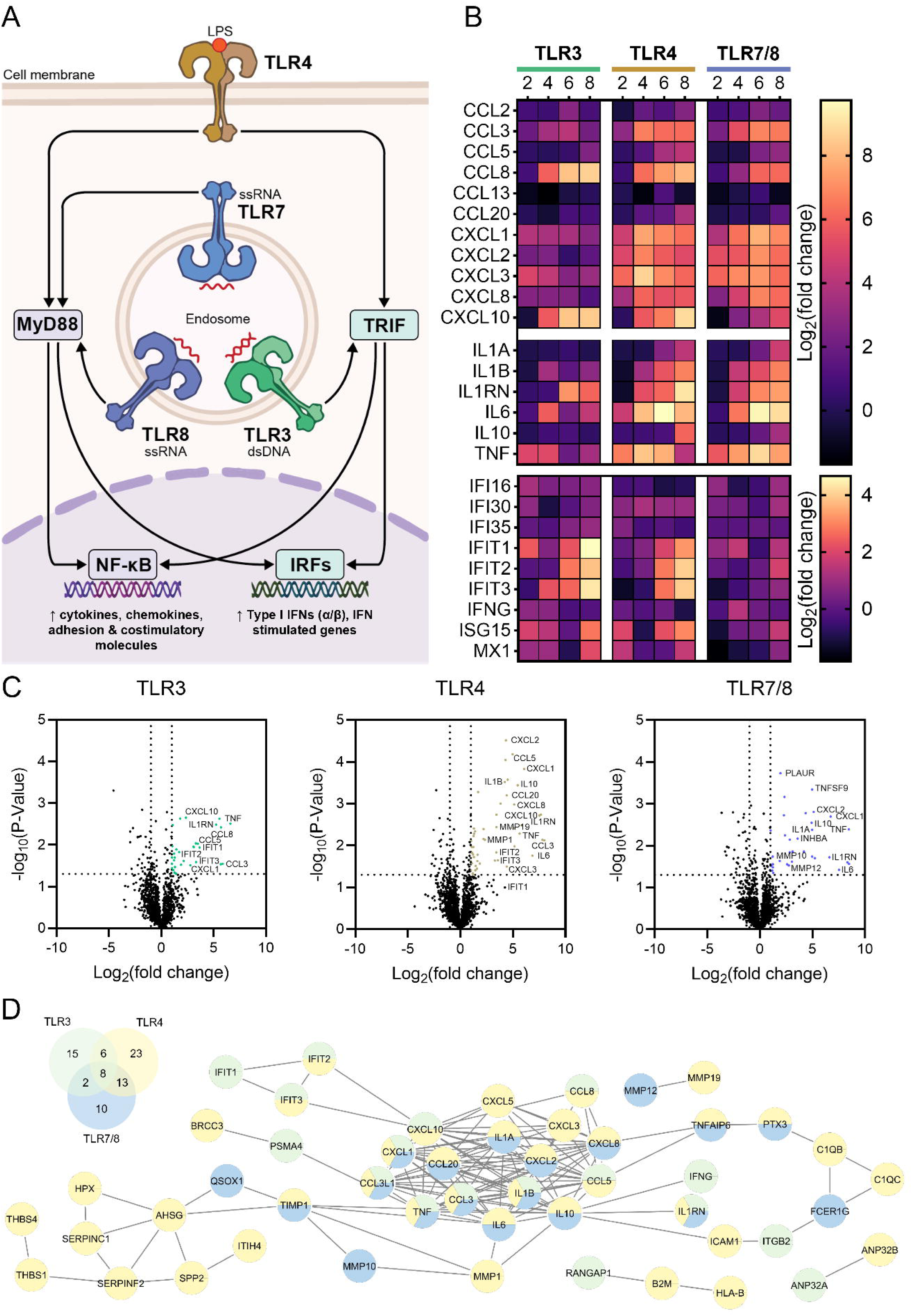
Profiling macrophage secretome responses to differential TLR activation. (A) Schematic overview of Toll-like receptor (TLR) signalling pathways activated by LPS (TLR4) and nucleic acid ligands (TLR3, TLR7/8), highlighting MyD88- and TRIF-dependent pathways leading to NF-κB activation and interferon regulatory factor (IRF) signalling, respectively. (B) Time-resolved log_2_(fold change) heatmap showing dynamic secretion of key cytokines, chemokines and interferon stimulated genes (ISGs) following TLR activation. Fold change values are calculated relative to corresponding M0 controls. (C) Volcano plots showing differential protein secretion at the 6-8 hour time point following TLR stimulation. Fold change values are calculated relative to corresponding M0 controls. (D) Combined venn diagram and protein-protein interaction network illustrating the overlap and connectivity of differentially secreted proteins relative to M0 controls across TLR3, TLR4 and TLR7/8 stimulation at the 6-8 hour time point. Nodes are coloured according to detection across stimulation conditions, with multi-coloured node indicating proteins shared between stimuli.

Expression of multiple Toll-like receptors, including TLR3, TLR4, TLR7 and TLR8, was first confirmed in resting iPSC-derived macrophages (Figure S4). Secretomic analysis of supernatants across all stimulation conditions and time points identified 1445 unique proteins (Table S5), including a broad array of pro-inflammatory cytokines, chemokines and ISGs. Focussing initially on these inflammatory mediators, temporal profiling revealed distinct, receptor-specific patterns of protein secretion (Figure 2B).

TLR3 activation elicited a pronounced interferon-driven response, consistent with activation of the TRIF-dependent signalling axis (45, 46), characterised by sustained release of ISGs, including C-X-C motif chemokine ligand 10 (CXCL10) and members of the interferon-induced protein with tetratricopeptide repeats (IFIT) family, from four hours onwards. In contrast, cytokine and chemokine secretion remained low under TLR3 stimulation relative to TLR4 and TLR7/8, with reduced secretion of key pro-inflammatory mediators such as tumour necrosis factor (TNF), interleukin-6 (IL-6) and multiple C-X-C motif chemokines, including CXCL8.

TLR4 stimulation produced a mixed secretion profile, consistent with engagement of both MyD88- and TRIF-dependent signalling pathways, resulting in robust secretion of pro-inflammatory chemokines and ISGs (47). Notably, the secretion of ISGs at the later time points is consistent with delayed TRIF-dependent signalling observed in response to LPS. Additionally, activation of this receptor was associated with increased secretion of the anti-inflammatory cytokine interleukin-10 (IL-10) at the 8-hour time point. In contrast, TLR7/8 activation resulted in a cytokine profile similar to TLR4, but with no secretion of ISGs, consistent with signalling predominantly through the MyD88 pathway (48).

To further characterise receptor-specific responses across the whole dataset, we focussed on the 6-8 hour time point, where differences between stimulation conditions were most clearly resolved. Differential expression analysis revealed both shared and receptor-specific patterns of protein secretion in response to each stimulus relative to M0 controls (Figure 2C). TLR4 stimulation resulted in a greater number of differentially secreted proteins and a higher overall magnitude of regulation compared to TLR3 and TLR7/8, consistent with its engagement of both MyD88- and TRIF-dependent signalling pathways. In contrast, TLR3 stimulation was enriched for ISGs, whereas TLR7/8 responses were dominated by pro-inflammatory cytokines and chemokines. Consistent with these observations, analysis of the corresponding intracellular proteomes demonstrated significant upregulation of interferon regulatory factors (IRFs) in response to TLR3 and TLR4 stimulation, but not TLR7/8 (Figure S5), supporting differential engagement of downstream signalling pathways.

A substantial degree of overlap was observed between stimulation conditions (Figure 2D), with a core set of pro-inflammatory cytokines, including interleukin-1β (IL-1β) and TNF, shared across all TLRs. Additionally, protein-protein interaction analysis highlighted the functional organisation of these responses (Figure 2D), revealing shared inflammatory modules alongside distinct clusters. Differential secretion of matrix metalloproteinases (MMPs) was evident between stimulation conditions, with MMP1 and MMP19 enriched following TLR4 activation, and MMP10 and MMP12 associated with TLR7/8 stimulation. These patterns suggest that distinct subsets of MMPs are engaged downstream of TLR4 and TLR7/8 activation, consistent with differential extracellular matrix remodelling processes linked to the resulting inflammatory responses (49).

Taken together, these results highlight the capacity of TLRs to drive both shared and specific secretory programmes, reflecting differential engagement of overlapping downstream signalling pathways. Importantly, our miniaturised sample preparation workflow coupled with dia-PASEF acquisition enables streamlined, quantitative profiling of macrophage responses through integrated proteomic and secretomics analysis, facilitating high-resolution interrogation of complex inflammatory networks.

### Development of a global concentration-response modelling pipeline

In drug discovery, concentration-response relationships describe how biological responses vary with compound concentration, providing a quantitative framework for assessing compound efficacy and safety (50). These analyses are typically applied to targeted assays, limiting the ability to capture broader, systems-level responses to drug perturbation. However, over the past decade, advances in mass spectrometry and data analysis workflows have enabled the extension of concentration-response analysis to proteome-wide datasets (50, 51). Recent studies have generated concentration-dependent profiles for thousands of proteins, enabling deeper characterisation of drug mechanisms of action and off-target effects (52, 53).

Building on these capabilities, we developed a global concentration-response modelling pipeline for DIA-based proteomic and secretomic datasets to quantify concentration-dependent effects on both intracellular protein abundance and secretion (Figure 3A). Concentration-response relationships were fitted using a four-parameter log-logistic function and compared to a null model to identify proteins exhibiting concentration-dependent behaviour (36). This approach was validated using proteomic and secretomic data from iPSC-derived macrophages from three human donors, in which TLR4 or TLR7/8 signalling was selectively inhibited across an 11-point concentration range. Fit quality was assessed using goodness-of-fit (R^2^), effect size and the differences in Akaike’s Information Criterion (ΔAIC) between the concentration-response and the null models. Proteins were classified as concentration-responsive using thresholds of R^2^ ≥ 0.5, to account for biological variability across donors, and an effect size of > 0.5 log units, to exclude flat or low-amplitude profiles that do not reflect biologically meaningful regulation. Additionally, calculated half maximum effective concentration (EC_50_) values were required to fall within the experimental concentration range (0.16 nM – 10 µM).

**Figure 3.**
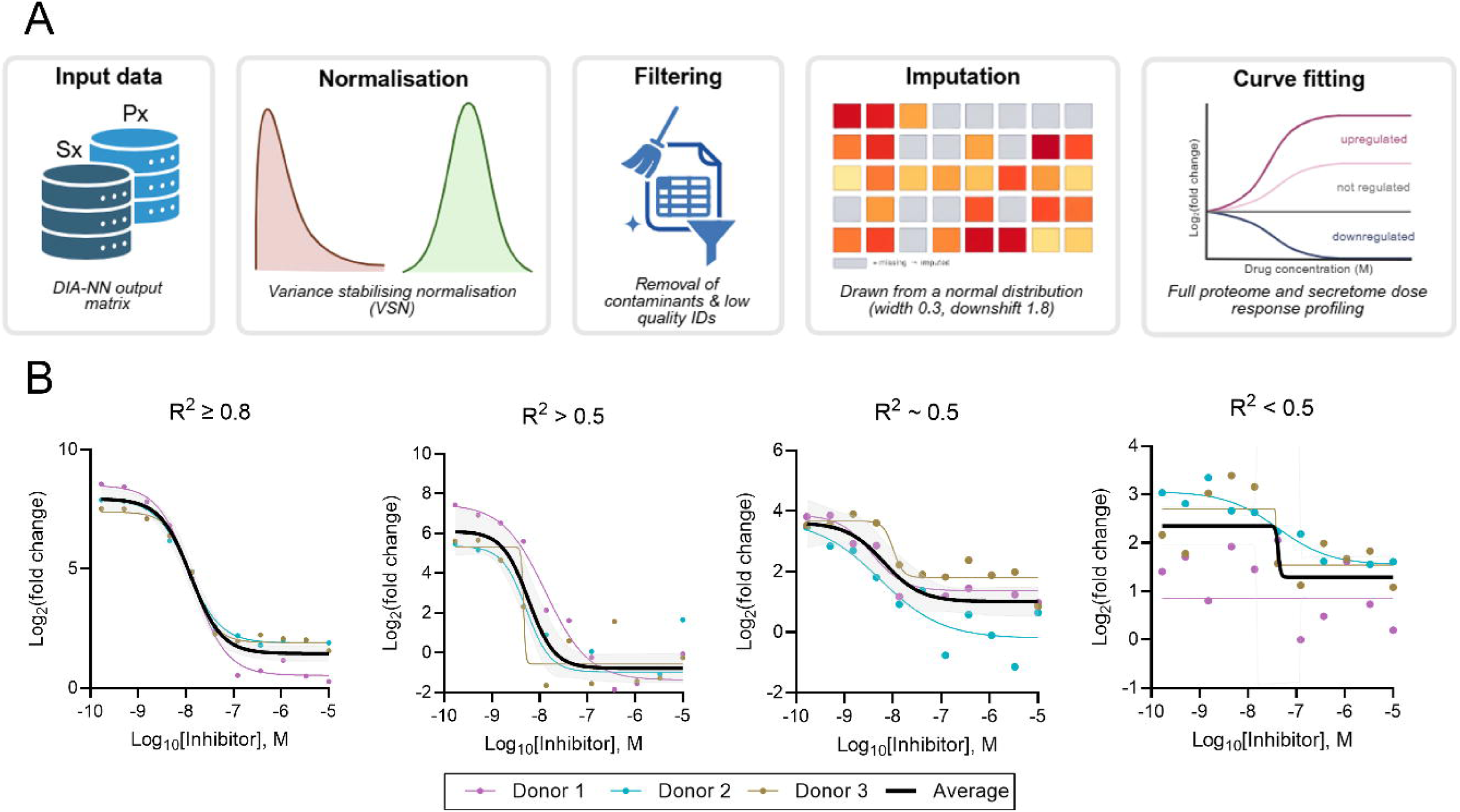
Data analysis pipeline for concentration-response curve fitting and fit performance assessment. (A) Schematic overview of the data analysis pipeline for proteome- and secretome-wide concentration-response analysis, including normalization, filtering of low-quality identifications (proteins lacking proteotypic peptides or present in less than two thirds of replicates), imputation of missing values and curve fitting. Concentration-response relationships were modelled using a four-parameter log-logistic function to quantify protein regulation relative to M0 controls. (B) Representative examples of curve fits spanning a range of goodness-of-fit values (R^2^), illustrating how fit quality distinguishes regulated from non-regulated proteins and supporting the use of an R^2^ cutoff of 0.5. Coloured curves represent fits to individual donors (n = 3), while the black curve represents the fit to combined data from all donors (n = 9). R² is calculated from the combined fit.

Representative curves spanning a range of R^2^ values illustrate how the fit quality criteria distinguishes well-defined concentration responsive behaviour from non-regulated profiles (Figure 3B), supporting the selected thresholds for classification. Having established this modelling framework, we next applied it to characterise the effects of TLR4 and TLR7/8 inhibition across the proteome and secretome of iPSC-derived macrophages.

### Global concentration-response profiling identifies receptor- and compartment-specific responses to TLR inhibition

To characterise global concentration-dependent effects of TLR pathway inhibition, iPSC-derived macrophages from three human donors were pre-treated with the selective TLR4 inhibitor resatorvid (TAK-242) or the TLR7/8 inhibitor MHV370 across an 11-point concentration range (0.16 nM – 10 µM) prior to receptor stimulation (54, 55). Matched cell lysate and supernatant fractions were collected for each condition and analysed using our miniaturised sample preparation workflow coupled with dia-PASEF acquisition to generate paired proteome and secretome datasets.

Concentration-response analysis of the secretomes following TLR4 and TLR7/8 inhibition identified 27 and 18 secreted proteins, respectively, meeting curve-fitting criteria (Tables S6 and S7; Figures S6 and S7). Substantial overlap was observed among cytokines and chemokines regulated by both inhibitors, including CXCL8 and TNF, which exhibited clear concentration-dependent reductions in secretion across the concentration range (Figure 4A). For these shared proteins, increasing inhibitor concentration progressively returned secretion levels towards those observed in unstimulated M0 controls (Figure 4A). In addition, 11 proteins were identified as uniquely concentration-responsive following TLR4 inhibition, comprising interferon-associated factors and matrix metalloproteinases previously highlighted in the time-resolved kinetics analysis. CXCL10 displayed a clear concentration-dependent reduction in secretion following TLR4 inhibition, whereas it did not exhibit robust concentration-responsive behaviour under TLR7/8 inhibition, despite being detected in that dataset (Figure 4A). In contrast, IL-12β was detected exclusively in the TLR7/8 secretome and exhibited concentration-dependent inhibition with increasing MHV370 concentration, in line with previously described TLR7/8-dependent secretion of this protein (55). Across concentration-responsive secreted proteins, pEC_50_ values were consistently higher for MHV370 (mean = 8.0) than for TAK-242 (mean = 6.3), indicating higher apparent potency under these conditions (Figure 4B).

**Figure 4.**
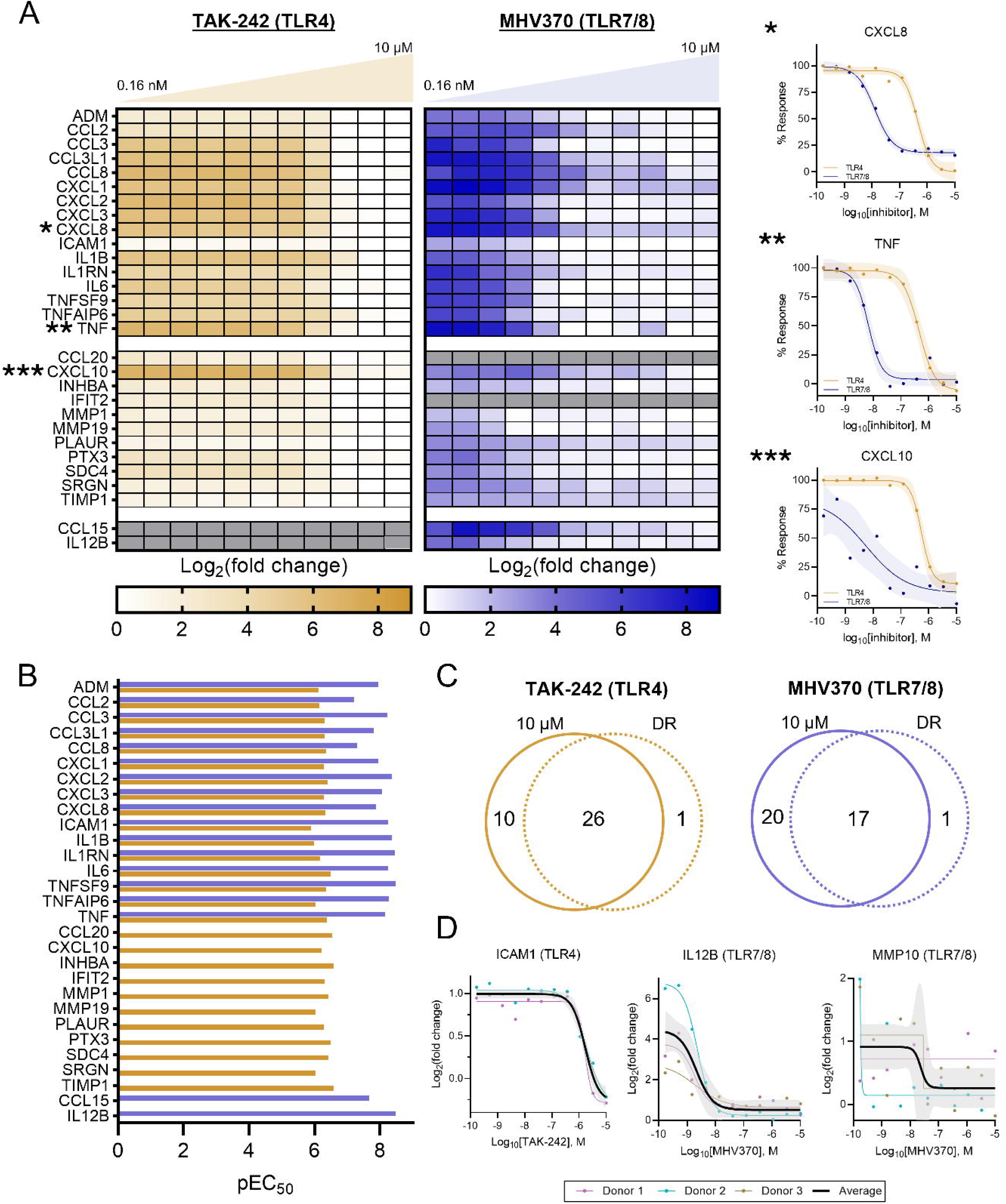
Concentration-response secretomics for profiling TLR pathway inhibition. (A) Heatmaps showing log_2_(fold change) of secreted protein abundance relative to M0 controls across an 11-point inhibitor concentration range (0.16 nM – 10 µM) for TAK-242 (TLR4) and MHV370 (TLR7/8). Proteins are grouped according to concentration-response fitting outcomes: those passing quality thresholds in both datasets (top), or selectively in the TLR4 (middle) or TLR7/8 (bottom) datasets. Three representative concentration-response curves (right) illustrate the range of fitted responses; curves represent the fit to combined data across donors (n = 9) with 95% confidence intervals. (B) Forest plot of pEC_50_ values derived from concentration-response curves for secreted proteins following treatment with TAK-242 (orange) and MHV370 (blue), illustrating inhibitor potencies across the secretome. (C) Venn diagrams summarizing the overlap between proteins identified as hits at the highest inhibitor concentration (10 µM) and those exhibiting concentration-responsive (DR) behaviour based on curve fitting criteria. (D) Representative curves highlighting differences between single-concentration hit identification and concentration-response clarification. ICAM1 and IL-12β exhibit concentration-responsive behaviour despite not being identified as statistically significant hits at 10 µM, while MMP10 is identified as a hit at 10 µM but does not display a clear concentration-response relationship.

To distinguish concentration-dependent protein secretion from effects observed only at a single concentration, we compared proteins meeting concentration-response criteria with those identified as significantly regulated at the highest concentration tested (10 µM). For both conditions, a substantial overlap was observed between concentration-responsive proteins and those identified as hits at 10 µM (Figure 4C). However, a substantial number of proteins identified as significantly regulated at the highest concentration did not pass curve fitting criteria (Table S8). Representative examples illustrate how concentration-response modelling resolves these distinctions (Figure 4D). ICAM1 and IL-12β both exhibited clear, concentration-responsive inhibition despite falling just below statistical significance at 10 µM, whereas MMP10 was identified as a significant hit in the TLR7/8 dataset at 10 µM but did not display a consistent concentration-response profile.

We next sought to compare concentration-responsive changes observed in the secretome with corresponding intracellular proteomic responses to assess overlap and relative potency across compartments. Concentration-response analysis of the intracellular proteomes following TLR4 and TLR7/8 inhibition identified 70 and 50 proteins, respectively, meeting curve-fitting criteria (Tables S9 and S10; Figures S8 and S9). A subset of proteins, largely consisting of pro-inflammatory cytokines and chemokines, were identified as concentration responsive in the proteome and secretome of both conditions (Figure 5A), with pEC_50_ values showing good agreement and most shared proteins falling within ± 0.5 log units. Apparent potency for most proteins was slightly higher in the proteome than the secretome, likely reflecting continued secretion of previously synthesised cytokines and chemokines and a delay between intracellular production and extracellular release.

**Figure 5.**
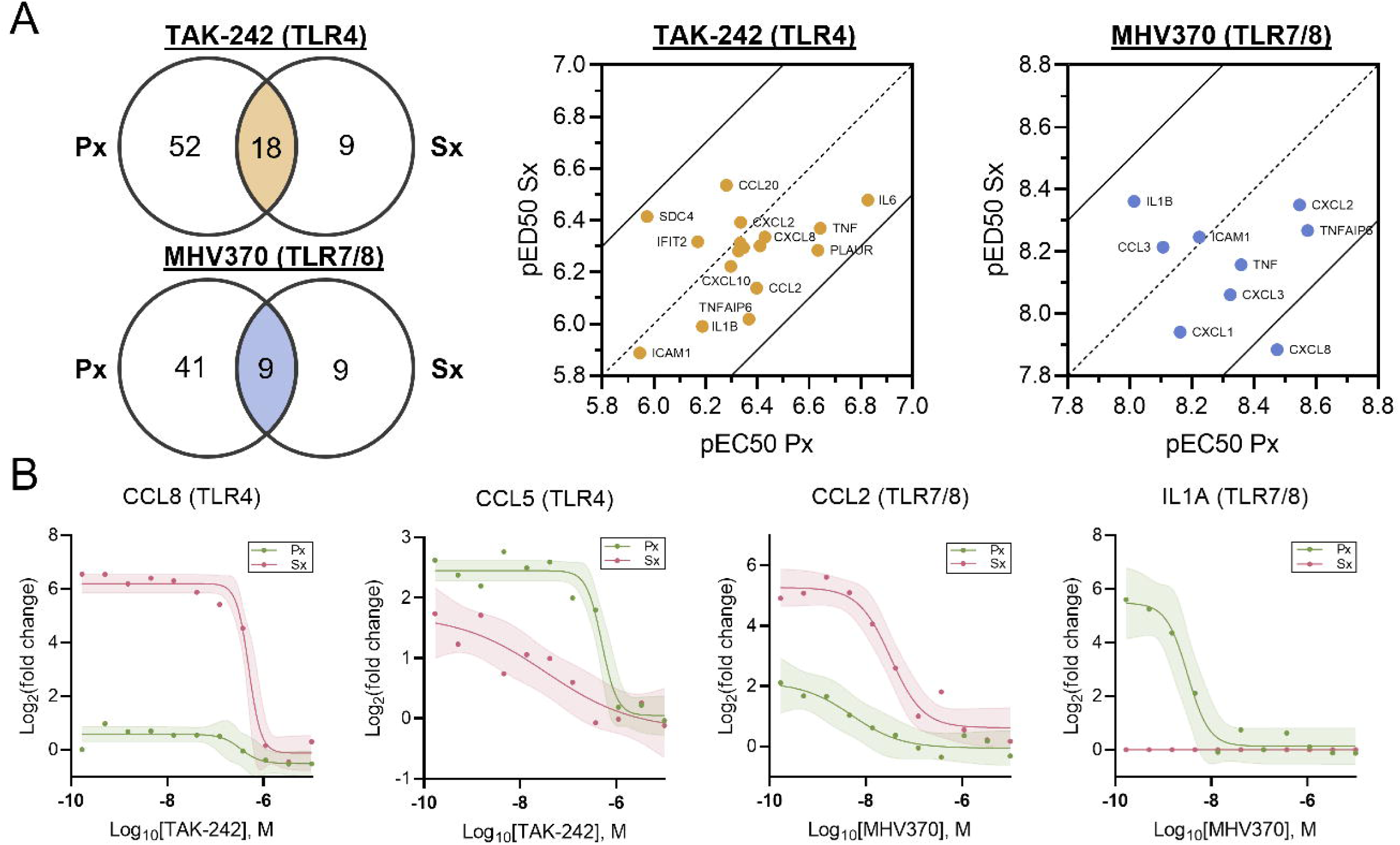
Comparative concentration-response analysis reveals pathway- and condition-specific inhibition of TLR4 and TLR7/8 signalling. (A) Venn diagrams showing the overlap of concentration-responsive proteins between the proteome (Px) and secretome (Sx) for TLR4 inhibition (TAK-242) and TLR7/8 inhibition (MHV370), highlighting shared and compartment-specific responses. Scatter plots of pEC50 values for proteins identified as concentration-responsive in both compartments demonstrate consistent inhibitor potency across targets, with values for each compartment falling within ± 0.5 log units. (B) Representative concentration-response curves for secreted cytokines and chemokines that exhibit differential expression profiles in the secretome compared to the proteome, following treatment with TAK-242 (TLR4; CCL8, CCL5) and MHV370 (TLR7/8; CCL2, IL-1α).

Additionally, a small number of cytokines and chemokines were uniquely identified as concentration-responsive in one compartment (Figure 5B). For example, CCL8 and CCL2 exhibited clear concentration-response behaviour in the secretome following TLR4 and TLR7/8 inhibition, respectively, whereas corresponding intracellular changes did not meet curve-fitting criteria. In contrast, CCL5 and IL-1α were identified as concentration-responsive only in the proteome, with CCL5 displaying a poorly defined response and IL-1α not detected in the secretome (Figure 5B). These compartment-specific differences highlight the distinct regulatory dynamics between intracellular protein abundance and extracellular secretion, including differences in protein synthesis, trafficking and secretion dynamics.

Collectively these results demonstrate that global, parallel concentration-response profiling across the proteome and secretome enables a comprehensive assessment of pharmacological perturbation of inflammatory signalling. By capturing concentration-dependent changes in both intracellular protein abundance and extracellular secretion, our approach resolves both pathway- and compartment-specific responses that would not be apparent from either dataset alone. This integrated approach provides a systems-level view of pathway modulation, enabling simultaneous evaluation of upstream regulatory effects and downstream functional outputs.

### Intracellular pathway responses to TLR4 and TLR7/8 inhibition

Finally, we examined intracellular concentration-responsive proteins to characterise pathway-level responses to TLR4 and TLR7/8 inhibition. Network and pathway enrichment analysis of proteins identified across both conditions revealed functional clustering into downstream signalling pathways (Figure 6A). Clusters shared between TLR4 and TLR7/8 inhibition were enriched for NF-κB signalling and cytokine-associated processes (Figure 6Aii), representing the core inflammatory responses mediated by MyD88-dependent signalling pathways (47). In contrast, TLR4-specific clusters were enriched for additional cytokine-associated proteins, including IL-6, alongside a distinct cluster corresponding to type I interferon signalling (Figure 6Ai), consistent with engagement of TRIF-dependent signalling pathways (47). Concentration-response analysis of these proteins, including ISG15 and IFIT2, demonstrated robust and well-defined inhibition with increasing TAK-242 concentration (Figure 6B), consistent with effective suppression of interferon-driven signalling at the intracellular level.

**Figure 6.**
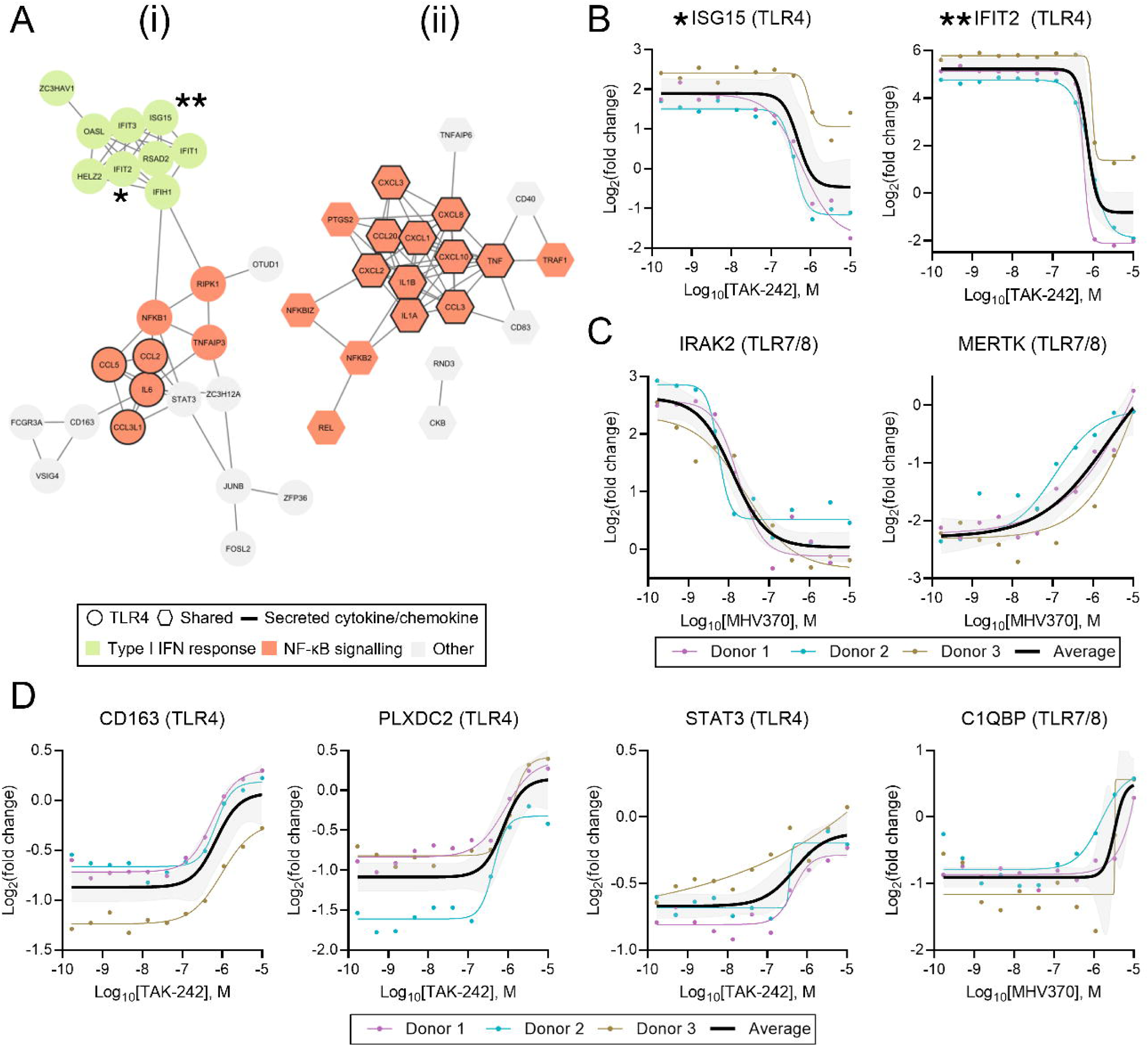
Intracellular pathway enrichment and concentration-dependent proteomic responses to TLR4 and TLR7/8 inhibition. (A) Network and pathway enrichment of concentration-responsive proteins from the proteomics (Px) datasets reveals functional clustering into key pathways. Nodes are coloured by pathway annotation and shaped to indicate TLR4-specific or shared regulation with TLR7/8. (B) Representative concentration-response curves for TLR4-specific interferon-related proteins following TAK-242 treatment, illustrating robust inhibitor-dependent suppression. (C) representative concentration-response curves for TLR7/8-specific proteins following MHV370 treatment, showing inhibitor-dependent changes in protein abundance. (D) Concentration-response curves for selected proteins (CD163, PLXDC2, STAT3 for TLR4; C1QBP for TLR7/8) that increase in abundance with inhibitor concentration, consistent with reversal towards the unstimulated (M0) phenotype.

Proteins identified as specifically concentration-responsive in the TLR7/8 dataset did not exhibit clear pathway-level clustering, but included proteins associated with inflammatory and TLR signalling pathways (Table S11). Representative examples include interleukin-1 receptor-associated kinase-like 2 (IRAK2), a direct downstream component of TLR signalling (56), which showed clear concentration-dependent inhibition, and Mer receptor tyrosine kinase (MERTK), a key regulator of efferocytosis and anti-inflammatory signalling (57), which exhibited increasing abundance with MHV370 concentration, consistent with restoration of a homeostatic, anti-inflammatory phenotype (Figure 6C).

Building on this observation, we identified multiple intracellular proteins that exhibited opposing concentration-dependent responses, with increasing abundance observed with inhibitor concentration and a return towards M0 baseline levels (Figure 6D). Representative examples include cluster of differentiation 163 (CD163), a macrophage scavenger receptor associated with anti-inflammatory responses (58); plexin domain containing protein 2 (PLXDC2), a critical cell-surface receptor that modulates macrophage polarisation and cytokine secretion (59); and signal transducer and activator of transcription 3 (STAT3), a transcription factor that is a key driver of anti-inflammatory (M2) macrophage polarisation (60), all of which increased in abundance following inhibition of TLR4 signalling. Similarly, complement component 1 Q subcomponent binding protein (C1QBP) exhibited the same trend following TLR7/8 inhibition, a multifunctional protein linked to mitochondrial function and the regulation of downstream inflammatory signalling (61). Together with the coordinated inhibition of pro-inflammatory processes, these trends highlight a coordinated shift towards baseline resting macrophage states following pharmacological inhibition of TLR signalling.

Collectively, these observations highlight the ability of this approach to capture both inhibition of pro-inflammatory signalling pathways and complementary responses that shift macrophages towards a resting or anti-inflammatory state, providing a comprehensive view of intracellular protein network regulation in response to pharmacological perturbation.

## Discussion

The human secretome represents a diverse set of proteins that are actively secreted or shed into the extracellular environment and play central roles in intercellular communication, immune regulation and tissue homeostasis (1, 2). As such, it provides a valuable resource for studying cellular responses to stimulation and pharmacological perturbation. Conventional immunoassays remain the gold standard for measuring secreted proteins but rely on predefined antibody panels and prior knowledge of targets, potentially limiting their ability to capture broader secretory responses (15–17). In contrast, MS-based secretomics approaches offer an unbiased alternative, with recent advances in acquisition strategies, including dia-PASEF, significantly improving throughput and proteomic depth without compromising sensitivity (27). However, their application remains constrained by sample preparation workflows that rely on protein precipitation, limiting reproducibility and scalability in drug discovery settings (28–30).

Here, we addressed this limitation by developing and validating a plate-based ISD-SPE workflow enabling miniaturised, automatable processing of conditioned media in a standard 96-well format. Benchmarking against conventional acetone precipitation demonstrated comparable proteomic coverage with improved quantitative reproducibility, reflected by a reduction in median protein %CV from 8.8% to 5.1%, and strong agreement in fold changes between methods (R^2^ = 0.81). Furthermore, the absence of edge effects, low missing value rates and high inter-plate correlation (Pearson r > 0.9) indicates consistent performance across plates and supports application to large-scale study designs. By eliminating precipitation, drying and resolubilisation steps, ISD-SPE reduces handling complexity and improves compatibility with automated workflows, establishing a robust and scalable approach for secretome sample preparation.

Application of the ISD-SPE workflow to time-resolved profiling of macrophage responses to TLR3, TLR4 and TLR7/8 stimulation revealed clear receptor-specific differences in secretory behaviour, consistent with the differential engagement of MyD88- and TRIF-dependent signalling pathways (62). TLR3 responses were characterised by an interferon-driven profile with limited cytokine secretion (62), whereas TLR7/8 stimulation produced a complementary response dominated by pro-inflammatory cytokines and chemokines (63). TLR4 combined features of both pathways, producing a robust secretory response that reflect its capacity to signal via both MyD88 and TRIF (64). Receptor-specific differences in MMP secretion further suggest distinct extracellular remodelling responses downstream of TLR activation. Given the emerging roles of MMPs not only in extracellular matrix turnover but also in regulating inflammatory signalling and immune cell behaviour (65–67), these observations may warrant further investigation into how TLR-dependent MMP programmes contribute to shaping functional immune responses.

We also applied concentration-response modelling to matched proteome and secretome datasets following TLR4 and TLR7/8 inhibition. Whilst proteome-wide concentration-response profiling has been applied to characterise drug mechanisms of action in cell lysates (50–52), its application to the secretome, particularly in the context of paired proteome and secretome data from the same experiment, remains limited. Extending this approach to extracellular measurements enabled quantitative characterisation of concentration-dependent effects on protein secretion, enabling the direct comparison of intracellular signalling events and downstream secretory outputs from the same sample well.

The use of a four-parameter log-logistic model allowed robust identification of proteins exhibiting consistent concentration-response behaviour across three human donors. Comparison with single-concentration analyses demonstrated that concentration-response modelling and single-concentration endpoints capture distinct aspects of pharmacological response. We found that proteins including ICAM1 and IL-12β displayed well-defined concentration-response profiles despite not reaching statistical significance at the highest concentration, whereas others such as MMP10, identified as significantly regulated at 10 µM, did not exhibit consistent concentration dependence. These observations highlight the added value of curve fitting in distinguishing reproducible pathway modulation from off-target or nonspecific effects observed at individual concentration points, and emphasise the benefit of integrating both analyses for confident identification of potential markers of pathway activity.

Integration of matched datasets enabled direct comparison of intracellular signalling and extracellular output. For proteins identified as concentration-responsive in both compartments, the overall agreement in pEC_50_ values supports a coupled relationship between pathway inhibition and downstream secretion. Direct comparison of pEC_50_ values further revealed that apparent potency was consistently higher in the proteome, reflecting a biological lag between intracellular inhibition and the corresponding extracellular response (68). The magnitude of this lag is likely influenced by protein-specific synthesis, processing and secretion dynamics.

Proteins exhibiting compartment-specific concentration-response behaviour provided further insight into these regulatory dynamics. For example, CCL8 and CCL2 showed clear concentration-responsive behaviour in the secretome, whereas the corresponding intracellular responses were less well defined, with weaker effect sizes and poorer curve fits that did not meet our concentration-response thresholds. As known secreted chemokines, both proteins are rapidly induced and released upon stimulation (69, 70), with the dominant extracellular signal reflecting this mode of regulation. In contrast, CCL5 and IL-1α were identified as concentration-responsive only in the proteome, with IL-1α undetected in corresponding secretomes. This is consistent with reports that CCL5 can be stored intracellularly and released in a regulated manner independent of de novo protein synthesis (71), as well as the predominantly intracellular localisation and context-dependent release of IL-1α (72). Together, these observations demonstrate that intracellular protein abundance and extracellular secretion do not always correlate, and that profiling both provides a more complete view of pathway modulation.

Analysis of intracellular concentration-responsive proteins provided additional pathway-level resolution of TLR signalling inhibition. The clear clustering of proteins regulated by TAK-242-mediated TLR4 inhibition into NF-κB and type I IFN signalling pathways confirms on-target pathway modulation (54). Notably, increasing inhibitor concentration across both TLR4 and TLR7/8 systems was also associated with restoration of proteins linked to homeostatic macrophage phenotypes, including CD163, PLXDC2 and STAT3. CD163 is a well-characterised cell surface marker of anti-inflammatory macrophage identity that is shed in response to pro-inflammatory stimuli such as LPS (58, 73), and its concentration-dependent recovery is consistent with restoration of a resting macrophage phenotype. Similarly, the observed increase in STAT3, a key mediator of IL-10-dependent anti-inflammatory signalling (74), provides an intracellular correlate to the IL-10 secretion observed at later time points in the time-resolved kinetics experiment. Together, these findings highlight that global concentration-response profiling captures not only the suppression of known pro-inflammatory signalling but also active restoration of homeostatic pathways, providing a more comprehensive assessment of compound effects.

In summary, this study establishes a scalable, plate-based secretomics workflow that enables integrated profiling of macrophage inflammatory responses across matched proteome and secretome datasets. The combination of ISD-SPE sample preparation, dia-PASEF acquisition and concentration-response modelling enables robust identification of concentration-dependent effects on both intracellular signalling and extracellular secretion of pro-inflammatory pathways and restoration of homeostatic processes, providing a more complete view of pharmacological modulation than either readout alone. Whilst demonstrated here using well-characterised TLR biology and a small number of tool compounds, our approach enables direct, system-wide assessment of cellular responses of pharmacological perturbation, with the potential to uncover new regulatory mechanisms and accelerate the evaluation of next-generation therapeutics.

## Data Availability

All MS raw files and output information from DIA-NN have been deposited in the ProteomeXchange Consortium via the PRIDE partner repository with the dataset identifier PXD081205. The source code used for concentration-response modelling is available on GitHub (https://github.com/lmcdbd/dose_response).

## Supplemental data

This article contains supplemental data.

## Acknowledgements

We would like to thank the Translational Models team at GSK for supporting cell culture, and the Sample Management team at GSK for the handling and dispensing of compound plates. This work was funded by GSK.

## Abbreviations

AIC: Akaike’s Information Criterion
CC: Cellular Component
CCL: C-C Motif Chemokine Ligand
C1QBP: Complement Component 1 Q Subcomponent Binding Protein
CV: Coefficient of Variation
CXCL: C-X-C Motif Chemokine Ligand
DIA: Data-Independent Acquisition
DIA-NN: Data-Independent Acquisition by Neural Networks
DPBS: Dulbecco’s Phosphate Buffered Saline
EC50: Effective Concentration at Half Maximum
ELISA: Enzyme-Linked Immunosorbent Assay
FBS: Fetal Bovine Serum
FDR: False Discovery Rate
GO: Gene Ontology
IAA: Iodoacetamide
IFIT: Interferon-Induced Protein with Tetratricopeptide Repeats
IFN: Interferon
IL: Interleukin
IRAK: Interleukin-1 Receptor-Associated Kinase
IRF: Interferon Regulatory Factor
ISD-SPE: In-Solution Digestion and Solid-Phase Extraction
ISG: Interferon-Stimulated Gene
LC-MS/MS: Liquid Chromatography Tandem Mass Spectrometry
LPS: Lipopolysaccharide
M0: Resting macrophage phenotype
M1: Pro-inflammatory macrophage phenotype
M-CSF: Macrophage Colony-Stimulating Factor
MERTK: MER Proto-Oncogene Tyrosine Kinase
MHV370: TLR7/8 inhibitor
MMP: Matrix Metalloproteinase
MS: Mass Spectrometry
MyD88: Myeloid Differentiation Primary Response Protein 88
nELISA: Nucleobase-Enabled Localised Immunoassay with Spectral Addressing
OptiMEM: Opti-MEM Reduced Serum Medium
PASEF: Parallel Accumulation Serial Fragmentation
PCA: Principal Component Analysis
PEA: Proximity Extension Assay
PLXDC2: Plexin Domain Containing Protein 2
PRIDE: Proteomics Identifications Database
PRR: Pattern Recognition Receptor
R2: Coefficient of Determination
R848: Resiquimod
RSS: Residual Sum of Squares
SDS: Sodium Dodecyl Sulphate
SPD: Samples Per Day
STAT3: Signal Transducer and Activator of Transcription 3
TAK-242: Resatorvid, TLR4 inhibitor
TCEP: Tris(2-carboxyethyl)phosphine
TEAB: Triethylammonium Acetate
TFA: Trifluoroacetic Acid
TIMS: Trapped Ion Mobility Spectrometry
TLR: Toll-Like Receptor
TNF: Tumour Necrosis Factor
TRIF: TIR-Domain-Containing Adapter Molecule 1
VSN: Variance Stabilising Normalisation

## References

1. Wu, W., and Krijgsveld, J. (2024) Secretome Analysis: Reading Cellular Sign Language to Understand Intercellular Communication. Mol. Cell. Proteomics 23, 10.1016/j.mcpro.2023.100692

2. Murray, R. Z., and Stow, J. L. (2014) Cytokine secretion in macrophages: SNAREs, Rabs, and membrane trafficking. Front. Immunol. 5, 10.3389/fimmu.2014.0053

3. Tjalsma, H., Antelmann, H., Jongbloed, J. D. H., Braun, P. G., Darmon, E., Dorenbos, R., Dubois, J. F., Westers, H., Zanen, G., Quax, W. J., Kuipers, O. P., Bron, S., Hecker, M., and Dijl, J. M. Van (2004) Proteomics of Protein Secretion by Bacillus subtilis: Separating the “Secrets” of the Secretome. Microbiol. Mol. Biol. Rev. 68, 207–233

4. Uhlén, M., Karlsson, M. J., Hober, A., Svensson, A., Scheffel, J., Kotol, D., Zhong, W., Tebani, A., Strandberg, L., Edfors, F., Sjöstedt, E., Mulder, J., Mardinoglu, A., Berling, A., Ekblad, S., Dannemeyer, M., Kanje, S., Rockberg, J., Lundqvist, M., Malm, M., Volk, A., Nilsson, P., Månberg, A., Dodig-crnkovic, T., Pin, E., Zwahlen, M., Oksvold, P., Feilitzen, K. Von, Häussler, R. S., Hong, M., Lindskog, C., Ponten, F., Katona, B., Vuu, J., Lindström, E., Nielsen, J., Robinson, J., Ayoglu, B., Mahdessian, D., Sullivan, D., Thul, P., Danielsson, F., Stadler, C., Lundberg, E., Bergström, G., Gummesson, A., Voldborg, B. G., Tegel, H., Hober, S., Forsström, B., Schwenk, J. M., Fagerberg, L., and Sivertsson, Å. (2019) The human secretome. Sci. Signal. 12, eaaz0274

5. Kaminska, P., Tempes, A., Scholz, E., and Malik, A. R. (2024) Cytokines on the way to secretion. Cytokine Growth Factor Rev. 79, 10.1016/j.cytogfr.2024.08.003

6. Lu, W., Zhang, Y., Ni, X., Wang, P., Zhuang, S., and Qin, W. (2025) Spatiotemporal profiling of modification-specific proteome secretion uncovers an itaconation-activated tyrosine kinase. Nat. Commun., 10.1038/s41467-025-66508-y

7. Duan, T., Du, Y., Xing, C., Wang, H. Y., and Wang, R. (2022) Toll-Like Receptor Signaling and Its Role in Cell-Mediated Immunity. Front. Immunol. 13, 10.3389/fimmu.2022.812774

8. Ding, M., Tegel, H., Sivertsson, Å., Hober, S., Snijder, A., Ormö, M., Strömstedt, P. E., Davies, R., and Holmberg Schiavone, L. (2020) Secretome-Based Screening in Target Discovery. SLAS Discov. 25, 535–551

9. Praznik, A., Fink, T., Franko, N., Lonzari, J., Ben, M., Ro, S., and Jerala, R. (2022) Regulation of protein secretion through chemical regulation of endoplasmic reticulum retention signal cleavage. Nat. Commun. 13, 10.1038/s41467-022-28971–9

10. Xue, H., Lu, B., and Lai, M. (2008) The cancer secretome: a reservoir of biomarkers. J. Transl. Med. 6, 10.1186/1479-5876-6–52

11. Meissner, F., Geddes-McAlister, J., Mann, M., and Bantscheff, M. (2022) The emerging role of mass spectrometry-based proteomics in drug discovery. Nat. Rev. Drug Discov. 21, 637–654

12. Evangelatos, G., Bamias, G., Kitas, G. D., Kollias, G., and Sfikakis, P. P. (2022) The second decade of antil1lTNFl1la therapy in clinical practice: new lessons and future directions in the COVIDl1l19 era Gerasimos. Rheumatol. Int. 42, 1493–1511

13. Rossi, M., and Young, J. W. (2005) Human Dendritic Cells: Potent Antigen-Presenting Cells at the Crossroads of Innate and Adaptive Immunity. J. Immunol. 175, 1373–1381

14. Broderick, L., Kambe, N., and Hoffman, H. M. (2026) Interleukin-1: from discovery to therapeutic targeting of the first cytokine. Allergol. Int. 75, 347–357

15. Belanger, L., Sylvestre, C., and Dufour, D. (1973) Enzyme-Linked Immunoassay for Alpha-Fetoprotein by Competitive and Sandwich Procedures. Clin. Chem. Acta 48, 15–18

16. Fu, Q., Zhu, J., and Van Eyk, J. E. (2010) Comparison of Multiplex Immunoassay Platforms. Clin. Chem. 56, 314–318

17. Chowdhury, F., Williams, A., and Johnson, P. (2009) Validation and comparison of two multiplex technologies, Luminex® and Mesoscale Discovery, for human cytokine profiling. J. Immunol. Methods 340, 55–64

18. Dagher, M., Ongo, G., Robichaud, N., Kong, J., Rho, W., Teahulos, I., Tavakoli, A., Bovaird, S., Merjaneh, S., Tan, A., Edwardson, K., Scheepers, C., Ng, A., Hajjar, A., Sow, B., Vrouvides, M., Lee, A., Decorwin-martin, P., Rasool, S., Huang, J., Erps, T., Coffin, S., Rashidi, N. M., Han, Y., Chandrasekaran, S. N., Miller, L., Kost-alimova, M., Skepner, A., Singh, S., Carpenter, A. E., Munzar, J. D., and Juncker, D. (2025) nELISA: a high-throughput, high-plex platform enables quantitative profiling of the inflammatory secretome. Nat. Methods 22, 2375–2385

19. Assarsson, E., Lundberg, M., Holmquist, G., Björkesten, J., Thorsen, S. B., Ekman, D., Eriksson, A., Dickens, E. R., Ohlsson, S., Edfeldt, G., Andersson, A. C., Lindstedt, P., Stenvang, J., Gullberg, M., and Fredriksson, S. (2014) Homogenous 96-plex PEA immunoassay exhibiting high sensitivity, specificity, and excellent scalability. PLoS One 9, 10.1371/journal.pone.0095192

20. Shuken, S. R. (2023) An Introduction to Mass Spectrometry-Based Proteomics. J. Proteome Res. 22, 2151–2171

21. Jiang, Y., Rex, D. A. B., Schuster, D., Neely, B. A., Rosano, G. L., Volkmar, N., Momenzadeh, A., Peters-Clarke, T. M., Egbert, S. B., Kreimer, S., Doud, E. H., Crook, O. M., Yadav, A. K., Vanuopadath, M., Hegeman, A. D., Mayta, M. L., Duboff, A. G., Riley, N. M., Moritz, R. L., and Meyer, J. G. (2024) Comprehensive Overview of Bottom-Up Proteomics Using Mass Spectrometry. ACS Meas. Sci. Au 4, 338–417

22. Kitata, R. B., Yang, J. C., and Chen, Y. J. (2022) Advances in data-independent acquisition mass spectrometry towards comprehensive digital proteome landscape. Mass Spectrom. Rev. e21781, 10.1002/mas.21781

23. Guo, T., Steen, J. A., and Mann, M. (2025) Mass-spectrometry-based proteomics: from single cells to clinical applications. Nature 638, 901–911

24. Demichev, V., Messner, C. B., Vernardis, S. I., Lilley, K. S., and Ralser, M. (2020) DIA-NN: neural networks and interference correction enable deep proteome coverage in high throughput. Nat. Methods 17, 41–44

25. Meier, F., Brunner, A. D., Koch, S., Koch, H., Lubeck, M., Krause, M., Goedecke, N., Decker, J., Kosinski, T., Park, M. A., Bache, N., Hoerning, O., Cox, J., Räther, O., and Mann, M. (2018) Online parallel accumulation–serial fragmentation (PASEF) with a novel trapped ion mobility mass spectrometer. Mol. Cell. Proteomics 17, 2534–2545

26. Meier, F., Park, M. A., and Mann, M. (2021) Trapped ion mobility spectrometry and parallel accumulation–serial fragmentation in proteomics. Mol. Cell. Proteomics 20, 10.1016/j.mcpro.2021.100138

27. Tayler, C. L., Bateman, S., Haslam, C., Müller, L., Norris, K., Rosa-roseberry, E., Martens, S., Yu, J., Dickinson, E., Booty, L., Beveridge, R., Rattray, N. J. W., Peltier-heap, R., Tayler, C. L., Bateman, S., Haslam, C., Müller, L., Norris, K., Rosa-roseberry, E., Martens, S., Yu, J., Dickinson, E., Booty, L., Beveridge, R., Rattray, N. J. W., and Peltier-heap, R. (2026) dia-PASEF Enables Rapid Profiling of the Human Secretome for Deeper Insights Into Cellular Authors dia-PASEF Enables Rapid Profiling of the Human Secretome for Deeper Insights Into Cellular Dynamics and Inflammatory Mechanisms. Mol Cell Proteomics 25, 10.1016/j.mcpro.2026.101597

28. Almeida-Marques, C., Rolfs, F., Piersma, S. R., Bijnsdorp, I. V., Pham, T. V., Knol, J. C., and Jimenez, C. R. (2024) Secretome processing for proteomics: A methods comparison. Proteomics 24, 10.1002/pmic.202300262

29. Jiang, L., He, L., and Fountoulakis, M. (2004) Comparison of protein precipitation methods for sample preparation prior to proteomic analysis. J. Chromatogr. A 1023, 317–320

30. Nickerson, J. L., and Doucette, A. A. (2020) Rapid and Quantitative Protein Precipitation for Proteome Analysis by Mass Spectrometry. J. Proteome Res. 19, 2035–2042

31. Armesilla-Diaz, A., Arenaz, M. P., Ashby, C., Blanco, D., D’Oria, E., Garuti, H., Gómez, V., González-Del-Río, R., Martínez-Hoyos, M., Meiler, E., Mendoza-Losana, A., Mohamet, L., Padrón-Barthe, L., Pérez, E., Pérez, L., Remuiñán, M. J., Rodríguez-Miquel, B., Segura-Carro, D., and Viera-Morilla, S. (2025) High-throughput screening of small molecules targeting Mycobacterium tuberculosis in human iPSC macrophages. Antimicrob. Agents Chemother. 69, 10.1128/aac.01613-24

32. Hailemariam, M., Eguez, R. V., Singh, H., Bekele, S., Ameni, G., Pieper, R., and Yu, Y. (2018) S-Trap, an Ultrafast Sample-Preparation Approach for Shotgun Proteomics. J. Proteome Res. 17, 2917–2924

33. Skowronek, P., Thielert, M., Voytik, E., Tanzer, M. C., Hansen, F. M., Willems, S., Karayel, O., Brunner, A. D., Meier, F., and Mann, M. (2022) Rapid and In-Depth Coverage of the (Phospho-) Proteome With Deep Libraries and Optimal Window Design for dia-PASEF. Mol. Cell. Proteomics 21, 100279

34. Huber, W., von Heydebreck, A., Sultmann, H., Poustka, A., and Vingron, M. (2002) Variance stabilization applied to microarray data calibration and to the quantification of differential expression. Bioinformatics 18, S96–S104

35. Ritchie, M. E., Phipson, B., Wu, D., Hu, Y., Law, C. W., Shi, W., and Smyth, G. K. (2015) limma powers differential expression analyses for RNA-sequencing and microarray studies. Nucleic Acids Res. 43, 10.1093/nar/gkv007

36. Ritz, C., Baty, F., Streibig, J. C., and Gerhard, D. (2015) Concentration-response analysis using R. PLoS One 10, 10.1371/journal.pone.0146021

37. Akaike, H. (1974) in IEEE Transactions on Automatic Control, pp 716–723.

38. Dennis, G., Sherman, B. T., Hosack, D. A., Yang, J., Gao, W., Lane, H. C., and Lempicki, R. A. (2003) DAVID: Database for Annotation, Visualization, and Integrated Discovery. Genome Biol. 4, 10.1186/gb-2003-4-9-r60

39. Szklarczyk, D., Kirsch, R., Koutrouli, M., Nastou, K., Mehryary, F., Hachilif, R., Gable, A. L., Fang, T., Doncheva, N. T., Pyysalo, S., Bork, P., Jensen, L. J., and Von Mering, C. (2023) The STRING database in 2023: protein-protein association networks and functional enrichment analyses for any sequenced genome of interest. Nucleic Acids Res. 51, D638–D646

40. Chen, R., Zou, J., Chen, J., Zhong, X., Kang, R., and Tang, D. (2025) Pattern recognition receptors: function, regulation and therapeutic potential. Signal Transduct. Target. Ther. 10, 10.1038/s41392-025-02264–1

41. El-zayat, S. R., and Sibaii, H. (2019) Toll-like receptors activation, signaling, and targeting: an overview. Bull. Natl. Res. Cent. 43:187, 10.1186/s42269-019-0227–2

42. Wesche, H., Henzel, W. J., Shillinglaw, W., Li, S., and Cao, Z. (1997) MyD88: An Adapter That Recruits IRAK to the IL-1 Receptor Complex. Immunity 7, 837–847

43. Yamamoto, M., Sato, S., and Hemmi, H. (2003) Role of Adaptor TRIF in the MyD88-Independent Toll-like Receptor Signaling Pathway. Science (80-.). 301, 640–643

44. Koppenol-Raab, M., Sjoelund, V., Manes, N. P., Gottschalk, R. A., Dutta, B., Benet, Z. L., Fraser, I. D. C., and Nita-Lazar, A. (2017) Proteome and secretome analysis reveals differential post-transcriptional regulation of Toll-like receptor responses. Mol. Cell. Proteomics 16, S172–S186

45. Hu, L., Cheng, Z., Chu, H., Wang, W., Jin, Y., and Yang, L. (2024) TRIF-dependent signaling and its role in liver diseases. Front. Cell Dev. Biol. 12, 10.3389/fcell.2024.1370042

46. Tachizaki, M., Sakamoto, S., Kobori, Y., Asano, Y., Kawaguchi, S., Seya, K., Tanaka, H., Morita, E., and Imaizumi, T. (2024) Interferon-stimulated gene 56 positively regulates Toll-like receptor 3-mediated CXCL10 expression in human renal proximal tubular epithelial cells. FEBS Open Bio 14, 1303–1319

47. Kawasaki, T., and Kawai, T. (2014) Toll-like receptor signaling pathways. Front. Immunol. 5, 10.3389/fimmu.2014.0046

48. Heil, F., Hemmi, H., Hochrein, H., Ampenberger, F., Kirschning, C., Akira, S., Lipford, G., Wagner, H., and Bauer, S. (2004) Species-Specific Recognition of Single-Stranded RNA via Till-like Receptor 7 and 8. Science (80-.). 303, 1526–1529

49. Huang, W. C., Sala-Newby, G. B., Susana, A., Johnson, J. L., and Newby, A. C. (2012) Classical macrophage activation up-regulates several matrix metalloproteinases through mitogen activated protein kinases and nuclear factor-κB. PLoS One 7,

50. Berner, N., Bayer, F. P., George, A., Kabella, N., Bergamasco, F. L., and Kuster, B. (2026) Proteome-wide concentration-response measurements for the characterization of drug mechanism of action. Targetome 2, 10.48130/targetome-0025–0011

51. Mateus, A., Kurzawa, N., Becher, I., Sridharan, S., Helm, D., Stein, F., Typas, A., and Savitski, M. M. (2020) Thermal proteome profiling for interrogating protein interactions. Mol. Syst. Biol. 16, 10.15252/msb.20199232

52. Zecha, J., Bayer, F. P., Wiechmann, S., Woortman, J., Berner, N., Müller, J., Schneider, A., Kramer, K., Abril-Gil, M., Hopf, T., Reichart, L., Chen, L., Hansen, F. M., Lechner, S., Samaras, P., Eckert, S., Lautenbacher, L., Reinecke, M., Hamood, F., Prokofeva, P., Vornholz, L., Falcomatà, C., Dorsch, M., Schröder, A., Venhuizen, A., Wilhelm, S., Médard, G., Stoehr, G., Ruland, J., Grüner, B. M., Saur, D., Buchner, M., Ruprecht, B., Hahne, H., The, M., Wilhelm, M., and Kuster, B. (2023) Decrypting drug actions and protein modifications by concentration- and time-resolved proteomics. Science (80-.). 380, 93–101

53. Eckert, S., Berner, N., Kramer, K., Schneider, A., Müller, J., Lechner, S., Brajkovic, S., Sakhteman, A., Graetz, C., Fackler, J., Dudek, M., Pfaffl, M. W., Knolle, P., Wilhelm, S., and Kuster, B. (2025) Decrypting the molecular basis of cellular drug phenotypes by concentration-resolved expression proteomics. Nat. Biotechnol. 43, 406–415

54. Matsunaga, N., Tsuchimori, N., Matsumoto, T., and Ii, M. (2011) TAK-242 (resatorvid), a small-molecule inhibitor of Toll-like receptor (TLR) 4 signaling, binds selectively to TLR4 and interferes with interactions between TLR4 and its adaptor molecules. Mol. Pharmacol. 79, 34–41

55. Hawtin, S., André, C., Collignon-Zipfel, G., Appenzeller, S., Bannert, B., Baumgartner, L., Beck, D., Betschart, C., Boulay, T., Brunner, H. I., Ceci, M., Deane, J., Feifel, R., Ferrero, E., Kyburz, D., Lafossas, F., Loetscher, P., Merz-Stoeckle, C., Michellys, P., Nuesslein-Hildesheim, B., Raulf, F., Rush, J. S., Ruzzante, G., Stein, T., Zaharevitz, S., Wieczorek, G., Siegel, R., Gergely, P., Shisha, T., and Junt, T. (2023) Preclinical characterization of the Toll-like receptor 7/8 antagonist MHV370 for lupus therapy. Cell Reports Med. 4, 10.1016/j.xcrm.2023.101036

56. Pereira, M., and Gazzinelli, R. T. (2023) Regulation of innate immune signaling by IRAK proteins. Front. Immunol. 14, 10.3389/fimmu.2023.1133354 OPEN

57. She, Y., Xu, X., Yu, Q., Yang, X., He, J., and Tang, X. X. (2023) Elevated expression of macrophage MERTK exhibits profibrotic effects and results in defective regulation of efferocytosis function in pulmonary fibrosis. Respir. Res. 24, 10.1186/s12931-023-02424–3

58. Skytthe, M. K., Graversen, J. H., and Moestrup, S. K. (2020) Targeting of cd163+ macrophages in inflammatory and malignant diseases. Int. J. Mol. Sci. 21, 10.3390/ijms21155497

59. Guan, Y., Du, Y., Wang, G., Gou, H., Xue, Y., Xu, J., Li, E., Chan, D. W., Wu, D., Xu, P., Ni, P., Xu, D., and Hu, Y. (2021) Overexpression of PLXDC2 in Stromal Cell-Associated M2 Macrophages Is Related to EMT and the Progression of Gastric Cancer. Front. Cell Dev. Biol. 9, 10.3389/fcell.2021.673295

60. Xia, T., Zhang, M., Lei, W., Yang, R., Fu, S., Fan, Z., Yang, Y., and Zhang, T. (2023) Advances in the role of STAT3 in macrophage polarization. Front. Immunol. 14, 10.3389/fimmu.2023.1160719

61. Wang, Q., Chai, D., Sobhani, N., Sun, N., Neeli, P., Zheng, J., and Tian, H. (2022) C1QBP regulates mitochondrial plasticity to impact tumor progression and antitumor immune response. Front. Physiol. 13, 10.3389/fphys.2022.1012112

62. Kawai, T., and Akira, S. (2010) The role of pattern-recognition receptors in innate immunity: Update on toll-like receptors. Nat. Immunol. 11, 373–384

63. O’Neill, L. A. J., and Bowie, A. G. (2007) The family of five: TIR-domain-containing adaptors in Toll-like receptor signalling. Nat. Rev. Immunol. 7, 353–364

64. Shen, H., Tesar, B. M., Walker, W. E., and Goldstein, D. R. (2008) Dual Signaling of MyD88 and TRIF Is Critical for Maximal TLR4-Induced Dendritic Cell Maturation. J. Immunol. 181, 1849–1858

65. Nissinen, L., and Kähäri, V. M. (2014) Matrix metalloproteinases in inflammation. Biochim. Biophys. Acta 1840, 2571–2580

66. Mukherjee, A., and Das, B. (2024) The role of inflammatory mediators and matrix metalloproteinases (MMPs) in the progression of osteoarthritis. Biomater. Biosyst. 13, 10.1016/j.bbiosy.2024.100090

67. Zamilpa, R., and Lindsey, M. L. (2010) Extracellular matrix turnover and signaling during cardiac remodeling following MI: Causes and consequences. J. Mol. Cell. Cardiol. 48, 558–563

68. Knecht, S., Eberl, H. C., Kreisz, N., Ugwu, U. J., Starikova, T., Kuster, B., and Wilhelm, S. (2023) An Introduction to Analytical Challenges, Approaches, and Applications in Mass Spectrometry-Based Secretomics. Mol. Cell. Proteomics 22, 10.1016/j.mcpro.2023.100636

69. Zlotnik, A., and Yoshie, O. (2012) The Chemokine Superfamily Revisited. Immunity 36, 705–716

70. Griffith, J. W., Sokol, C. L., and Luster, A. D. (2014) Chemokines and chemokine receptors: Positioning cells for host defense and immunity. Annu. Rev. Immunol. 32, 659–702

71. Persaud, A. T., Bennett, S. A., Thaya, L., Burnie, J., and Guzzo, C. (2022) Human monocytes store and secrete preformed CCL5, independent of de novo protein synthesis. J. Leukoc. Biol. 111, 573–583

72. Dinarello, C. A. (2018) Introduction to the interleukin-1 family of cytokines and receptors: Drivers of innate inflammation and acquired immunity. Immunol. Rev. 281, 5–7

73. Etzerodt, A., and Moestrup, S. K. (2013) CD163 and inflammation: Biological, diagnostic, and therapeutic aspects. Antioxidants Redox Signal. 18, 2352–2363

74. Hutchins, A. P., Diez, D., and Miranda-Saavedra, D. (2013) The IL-10/STAT3-mediated anti-inflammatory response: Recent developments and future challenges. Brief. Funct. Genomics 12, 489–498

