## Supplementary_Figures_Data for "Plate-based ISD-SPE enables dual proteome-secretome concentration-response profiling of TLR signalling in iPSC-derived macrophages"

### Title

### Institutions

[1] GSK, Stevenage, UK

[2] Newcastle University Bioscience Institute, Faculty of Medical Sciences, Newcastle Upon Tyne, UK

[3] PerkinElmer, Stevenage, UK

[4] University of Strathclyde Department for Pure and Applied Chemistry, Glasgow, UK

[5] University of Strathclyde Institute of Pharmacy and Biomedical Sciences, Glasgow, UK

**Supplemental Figures and Figure Legends S1 – S9**

**Supplemental Files**

This article contains supplemental data that are included as separate data files that contain the following information:

***Supplemental Table 0.*** Column descriptors for each subsequent supplemental table.

***Supplemental Table 1.*** Signals Image Artist (SIMA) analysis parameters.

***Supplemental Table 2.*** Summary of optimised dia-PASEF method.

***Supplemental Table 3.*** List of proteins uniquely identified in acetone precipitation and ISD-SPE workflows.

***Supplemental Table 4.*** List of proteins identified as outliers from the Bland-Altman systematic bias analysis.

***Supplemental Table 5.*** Summary of the interval-based secretome data from Poly(I:C), LPS and R848 treated iPSC-derived macrophages.

***Supplemental Table 6.*** Concentration-response curve fitting outputs derived from secretome profiling of TAK-242 treated iPSC-derived macrophages.

***Supplemental Table 7.*** Concentration-response curve fitting outputs derived from secretome profiling of MHV370 treated iPSC-derived macrophages.

***Supplemental Table 8.*** Summary of proteins identified as concentration-responsive versus those detected at the highest inhibitor concentration for both compounds.

***Supplemental Table 9.*** Concentration-response curve fitting outputs derived from intracellular proteome profiling of TAK-242 treated iPSC-derived macrophages.

***Supplemental Table 10.*** Concentration-response curve fitting outputs derived from intracellular proteome profiling of MHV370 treated iPSC-derived macrophages.

***Supplemental Table 11.*** Summary of proteins uniquely identified as concentration-responsive in the intracellular proteomes of MHV370 treated iPSC-derived macrophages.

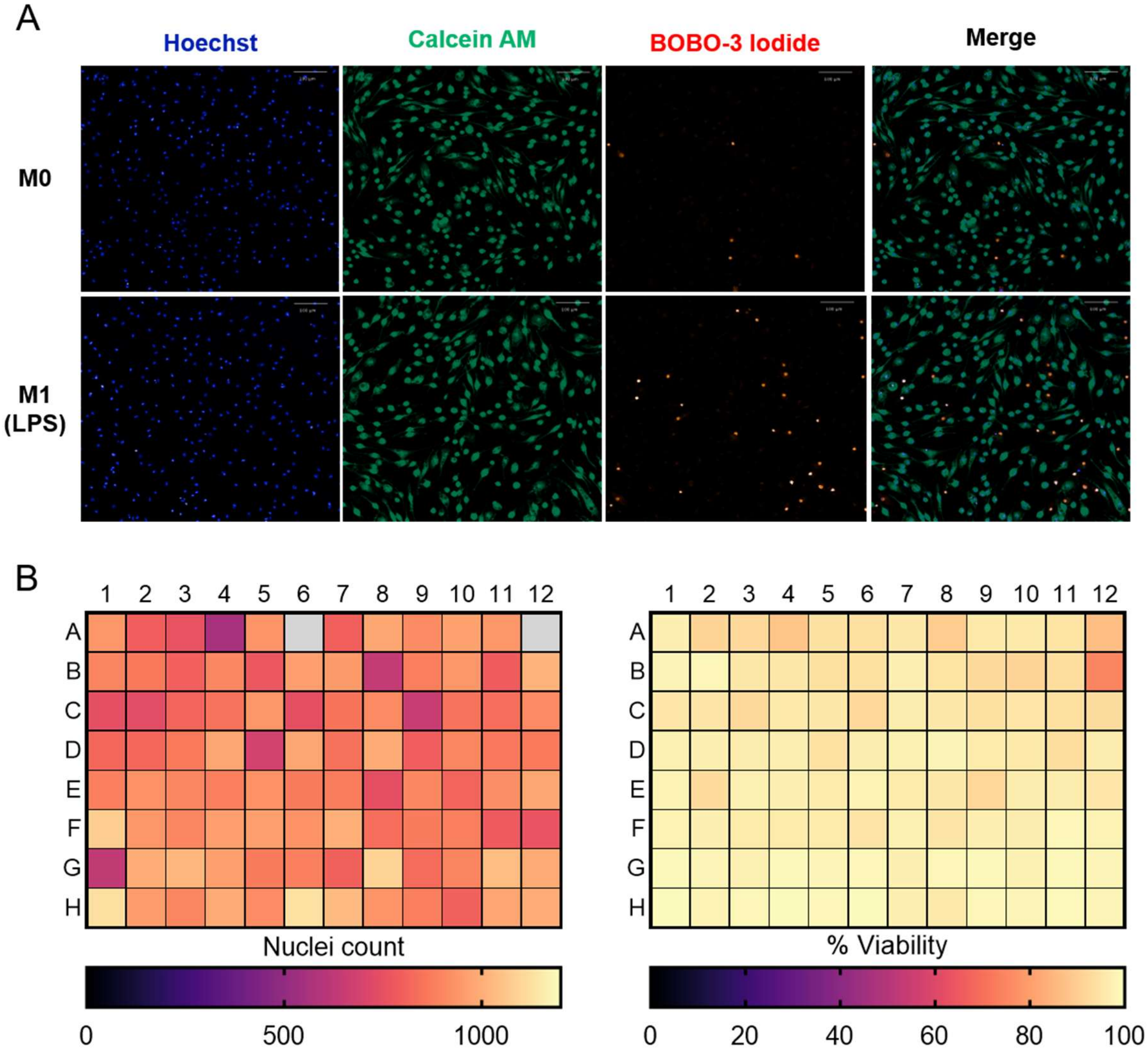


**Supplemental Fig. S1** Cell seeding uniformity and viability assessment of iPSC-derived macrophages under reduced serum conditions. (A) Fluorescence microscopy images show resting (M0) and LPS-stimulated (M1) macrophages after 4 hours in Opti-MEM™. Nuclei were stained with Hoechst (blue), viable cells with Calcein AM (green), and non-viable cells with BOBO-3 iodide (red). (B) Nuclei count (left) and cell viability (right) measurements for cells across the 96-well plate following 4 hours in Opti-MEM™, demonstrating uniform cell distribution and consistently high viability (mean = 95.6%)

under both M0 and M1 conditions. Grey wells indicate samples that were excluded from analysis due to a staining artefact.

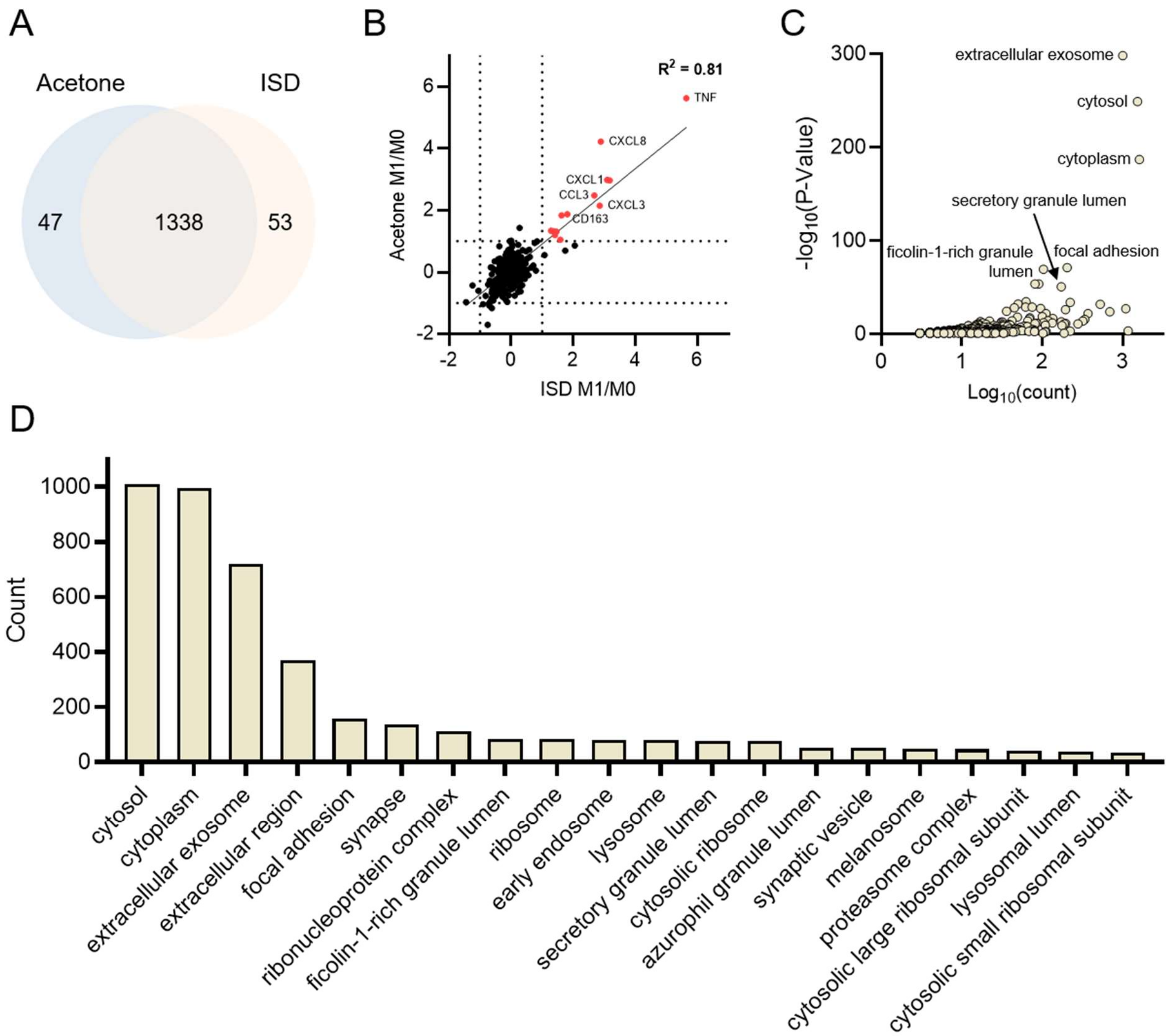


**Supplemental Fig. S2** Comparison between acetone precipitation and ISD-SPE workflows. (A) Venn diagram illustrates protein identification between acetone and ISD-SPE datasets. (B) $Log_2$(fold change) comparison between the two methods; fold changes calculated relative to M0 controls and filtered for $p < 0.05$. (C) Gene Ontology Cellular Component (GO CC) enrichment analysis showing extracellular and cytosolic protein annotations. (D) Number of proteins annotated to the top 20 significantly enriched GO CC terms identified in the ISD-SPE secretome dataset.

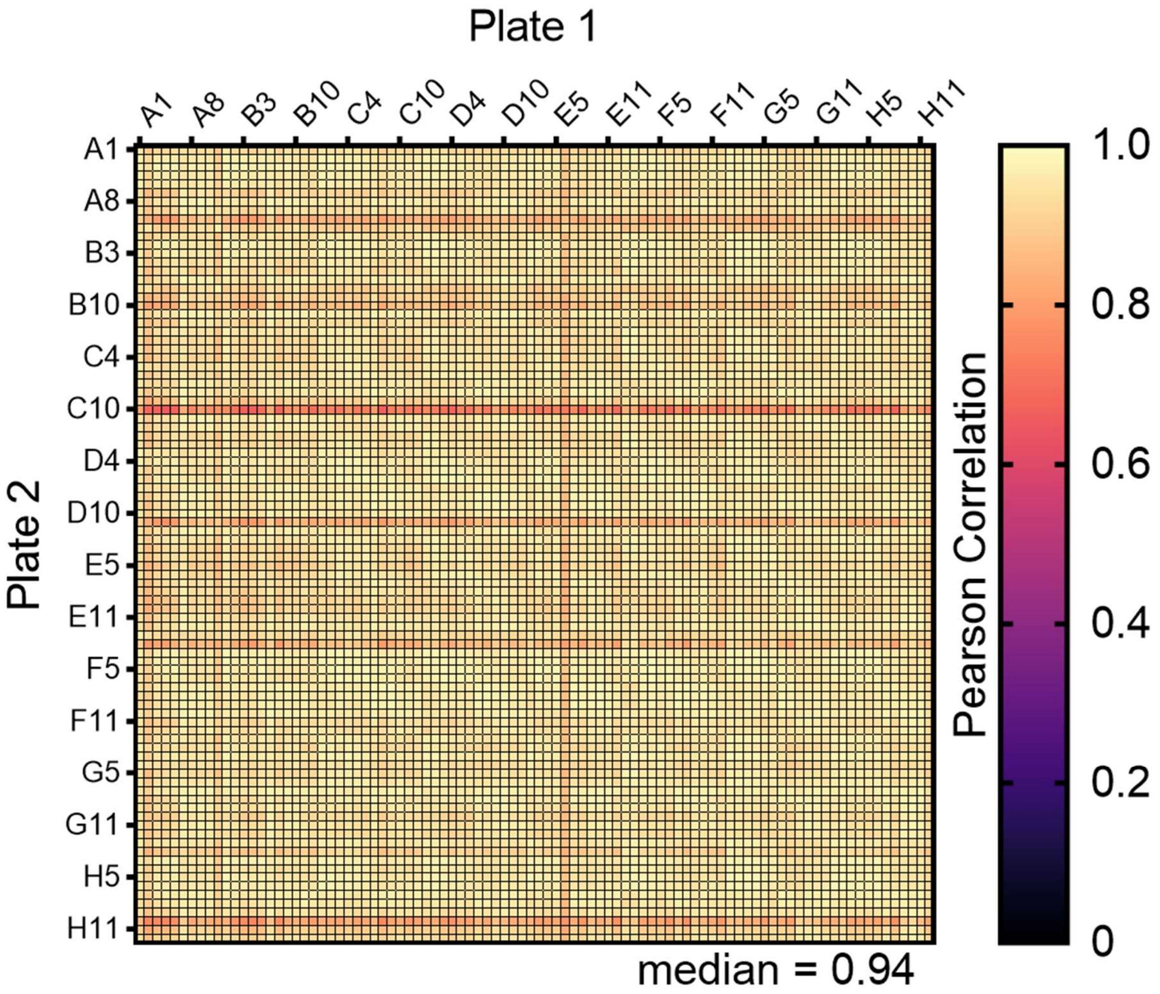


**Supplemental Fig. S3** Pairwise Pearson correlation analysis of protein abundances between two 96-well plates. Each cell represents the correlation between corresponding wells across plates, demonstrating strong overall agreement with a median correlation coefficient of 0.94.

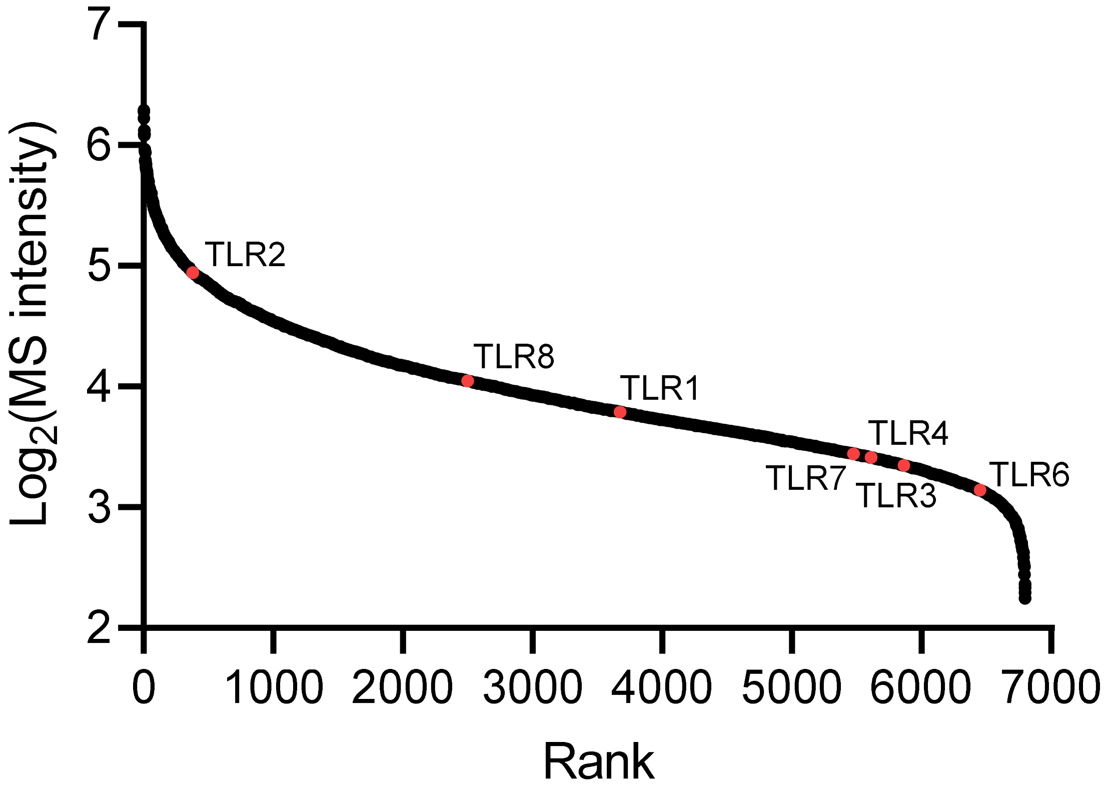


**Figure S4.** Detection of Toll-like receptors in resting iPSC-derived macrophages. Proteins are ranked by $log_2$-transformed MS intensity with Toll-like receptors highlighted in red.

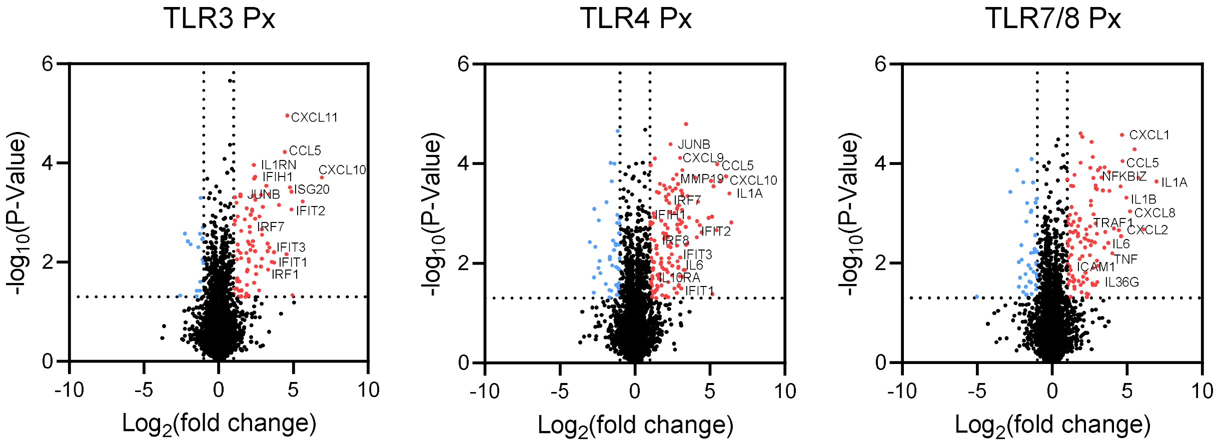


**Supplemental Fig. S5** Intracellular proteome responses to differential TLR activation after 8 hours. Volcano plots show differential protein abundance in the intracellular proteome (Px) following stimulation of TLR3, TLR4 and TLR7/8 for 8 hours, relative to M0 controls. $Log_2$(fold change) is plotted against $-log_{10}$(p-value). Significantly regulated proteins ($p < 0.05$ and fold change greater than 1) are highlighted, with selected interferon-stimulated genes (ISGs), cytokine-associated proteins and key transcriptional regulators annotated. The data highlights distinct yet overlapping intracellular responses to TLR activation, including enrichment of interferon signalling components for TLR3 and TLR4, consistent with differential engagement of TRIF-dependent pathways.

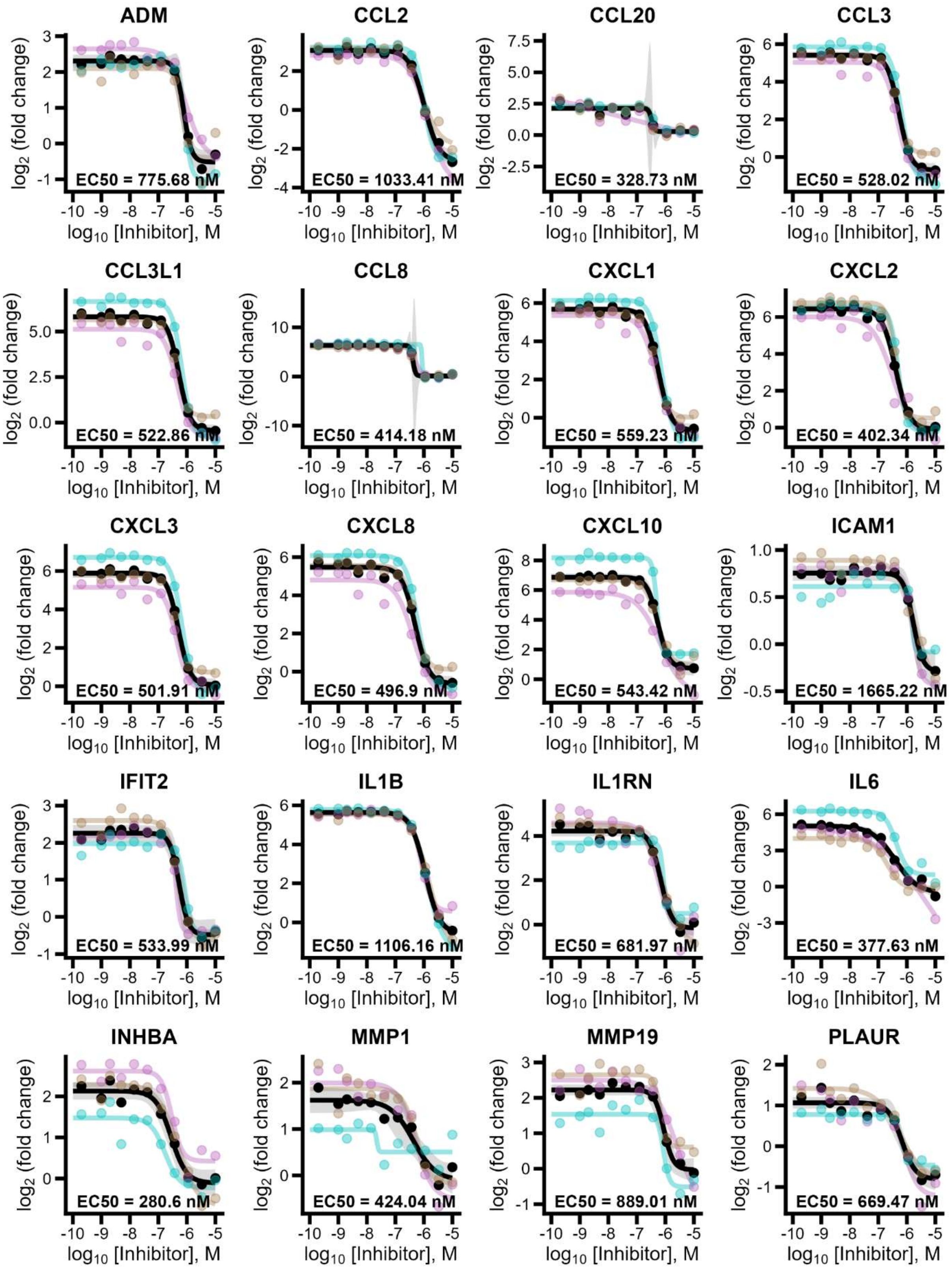
ADM
EC50 = 775.68 nM
CCL2
EC50 = 1033.41 nM
CCL20
EC50 = 328.73 nM
CCL3
EC50 = 528.02 nM
CCL3L1
EC50 = 522.86 nM
CCL8
EC50 = 414.18 nM
CXCL1
EC50 = 559.23 nM
CXCL2
EC50 = 402.34 nM
CXCL3
EC50 = 501.91 nM
CXCL8
EC50 = 496.9 nM
CXCL10
EC50 = 543.42 nM
ICAM1
EC50 = 1665.22 nM
IFIT2
EC50 = 533.99 nM
IL1B
EC50 = 1106.16 nM
IL1RN
EC50 = 681.97 nM
IL6
EC50 = 377.63 nM
INHBA
EC50 = 280.6 nM
MMP1
EC50 = 424.04 nM
MMP19
EC50 = 889.01 nM
PLAUR
EC50 = 669.47 nM
$log_2$ (fold change)
$log_{10}$ [Inhibitor], M

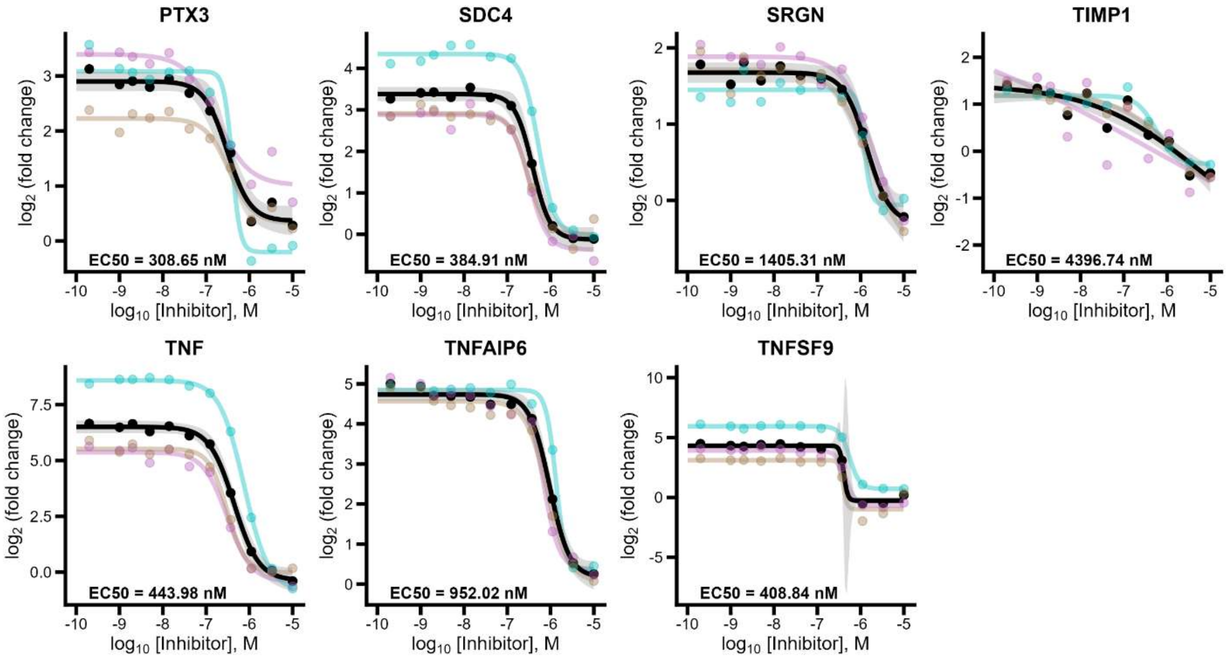


**Supplemental Fig. S6** Concentration-response curves showing $\log_2$(fold change) in secreted protein abundance relative to M0 controls across an 11-point inhibitor concentration range (0.16 nM – 10 µM) for TAK-242 (TLR4 inhibitor). Coloured curves represent fits to individual donors (n = 3), with corresponding data points shown, while the black curve represents the fit to combined data across donors (n = 9). Shaded regions indicate 95% confidence intervals. $EC_{50}$ values calculated from the combined fit are indicated for each protein. Selected proteins passed curve-fitting criteria and exhibit concentration-dependent suppression following TLR4 inhibition.

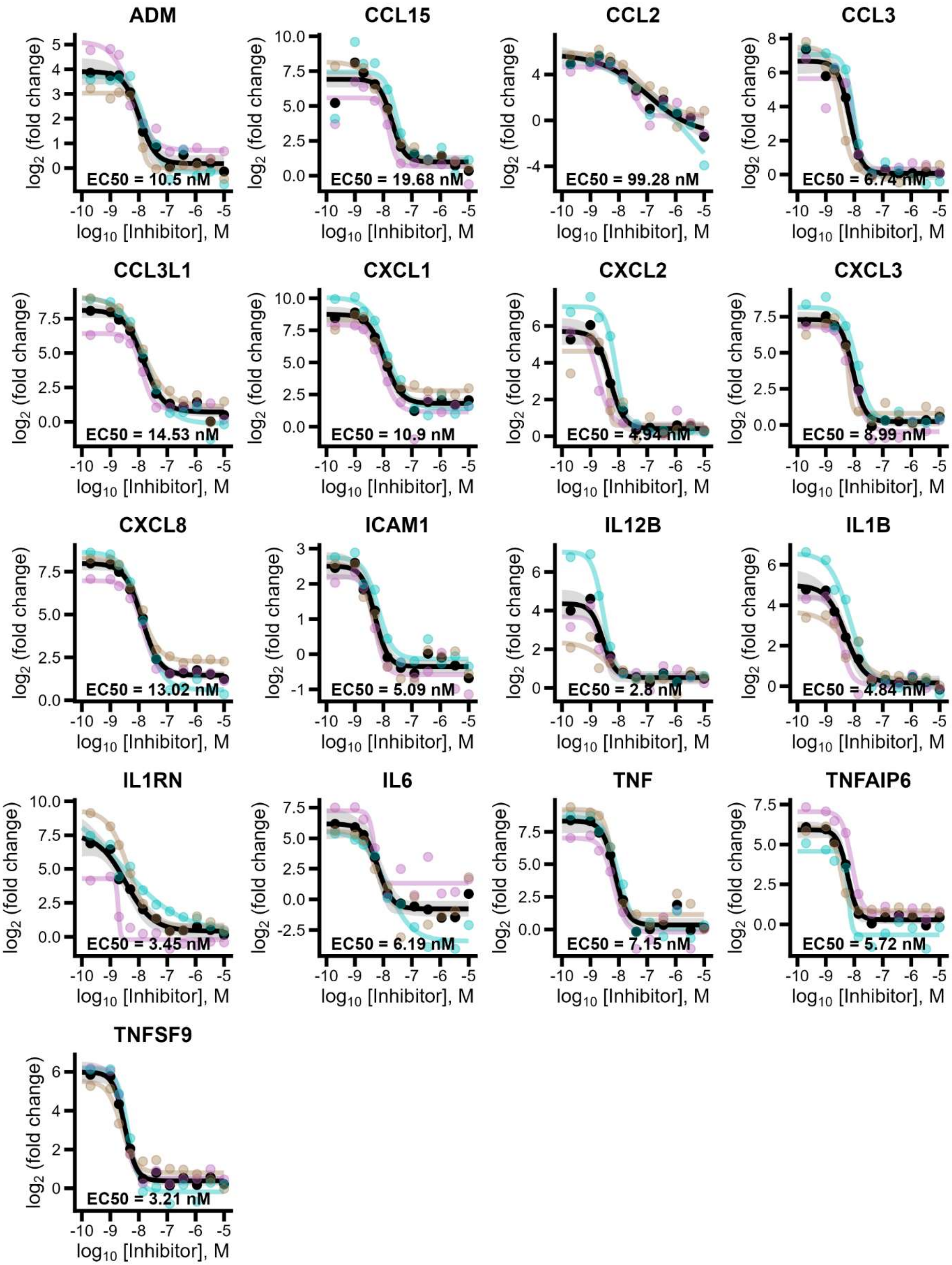
ADM
EC50 = 10.5 nM
CCL15
EC50 = 19.68 nM
CCL2
EC50 = 99.28 nM
CCL3
EC50 = 6.74 nM
CCL3L1
EC50 = 14.53 nM
CXCL1
EC50 = 10.9 nM
CXCL2
EC50 = 4.94 nM
CXCL3
EC50 = 8.99 nM
CXCL8
EC50 = 13.02 nM
ICAM1
EC50 = 5.09 nM
IL12B
EC50 = 2.8 nM
IL1B
EC50 = 4.84 nM
IL1RN
EC50 = 3.45 nM
IL6
EC50 = 6.19 nM
TNF
EC50 = 7.15 nM
TNFAIP6
EC50 = 5.72 nM
TNFSF9
EC50 = 3.21 nM
log2 (fold change)
log10 [Inhibitor], M

**Supplemental Fig. S7** Concentration-response curves showing $log_2$(fold change) in secreted protein abundance relative to M0 controls across an 11-point inhibitor concentration range (0.16 nM – 10 µM) for MHV370 (TLR7/8 inhibitor) Coloured curves represent fits to individual donors (n = 3), with corresponding data points shown, while the black curve represents the fit to combined data across donors (n = 9). Shaded regions indicate 95% confidence intervals. $EC_{50}$ values calculated from the combined fit are indicated for each protein. Selected proteins passed curve-fitting criteria and exhibit concentration-dependent suppression following TLR7/8 inhibition.

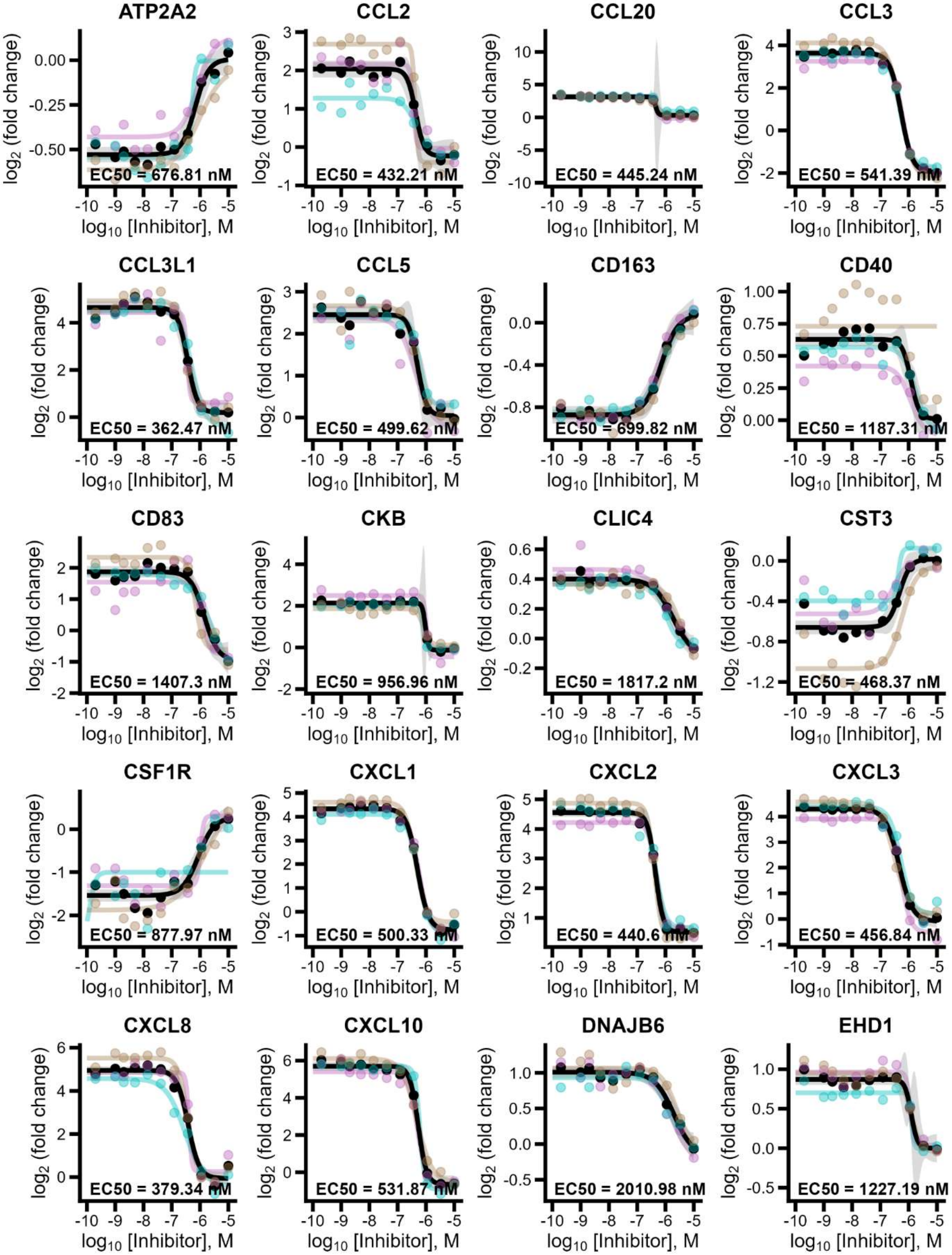
ATP2A2
EC50 = 676.81 nM
CCL2
EC50 = 432.21 nM
CCL20
EC50 = 445.24 nM
CCL3
EC50 = 541.39 nM
CCL3L1
EC50 = 362.47 nM
CCL5
EC50 = 499.62 nM
CD163
EC50 = 699.82 nM
CD40
EC50 = 1187.31 nM
CD83
EC50 = 1407.3 nM
CKB
EC50 = 956.96 nM
CLIC4
EC50 = 1817.2 nM
CST3
EC50 = 468.37 nM
CSF1R
EC50 = 877.97 nM
CXCL1
EC50 = 500.33 nM
CXCL2
EC50 = 440.6 nM
CXCL3
EC50 = 456.84 nM
CXCL8
EC50 = 379.34 nM
CXCL10
EC50 = 531.87 nM
DNAJB6
EC50 = 2010.98 nM
EHD1
EC50 = 1227.19 nM
log2 (fold change)
log10 [Inhibitor], M

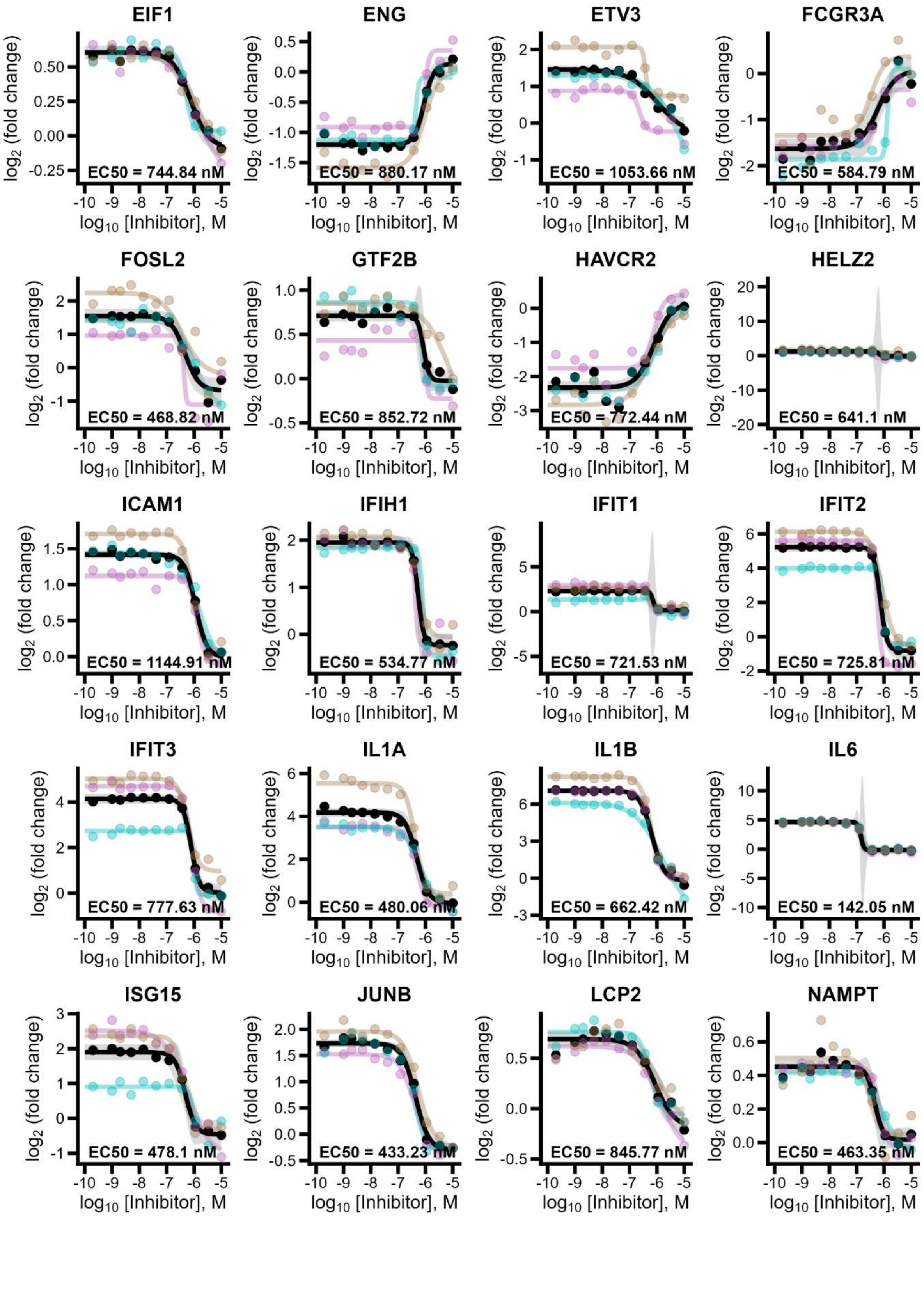
EIF1
EC50 = 744.84 nM
ENG
EC50 = 880.17 nM
ETV3
EC50 = 1053.66 nM
FCGR3A
EC50 = 584.79 nM
FOSL2
EC50 = 468.82 nM
GTF2B
EC50 = 852.72 nM
HAVCR2
EC50 = 772.44 nM
HELZ2
EC50 = 641.1 nM
ICAM1
EC50 = 1144.91 nM
IFIH1
EC50 = 534.77 nM
IFIT1
EC50 = 721.53 nM
IFIT2
EC50 = 725.81 nM
IFIT3
EC50 = 777.63 nM
IL1A
EC50 = 480.06 nM
IL1B
EC50 = 662.42 nM
IL6
EC50 = 142.05 nM
ISG15
EC50 = 478.1 nM
JUNB
EC50 = 433.23 nM
LCP2
EC50 = 845.77 nM
NAMPT
EC50 = 463.35 nM
log2 (fold change)
log10 [Inhibitor], M

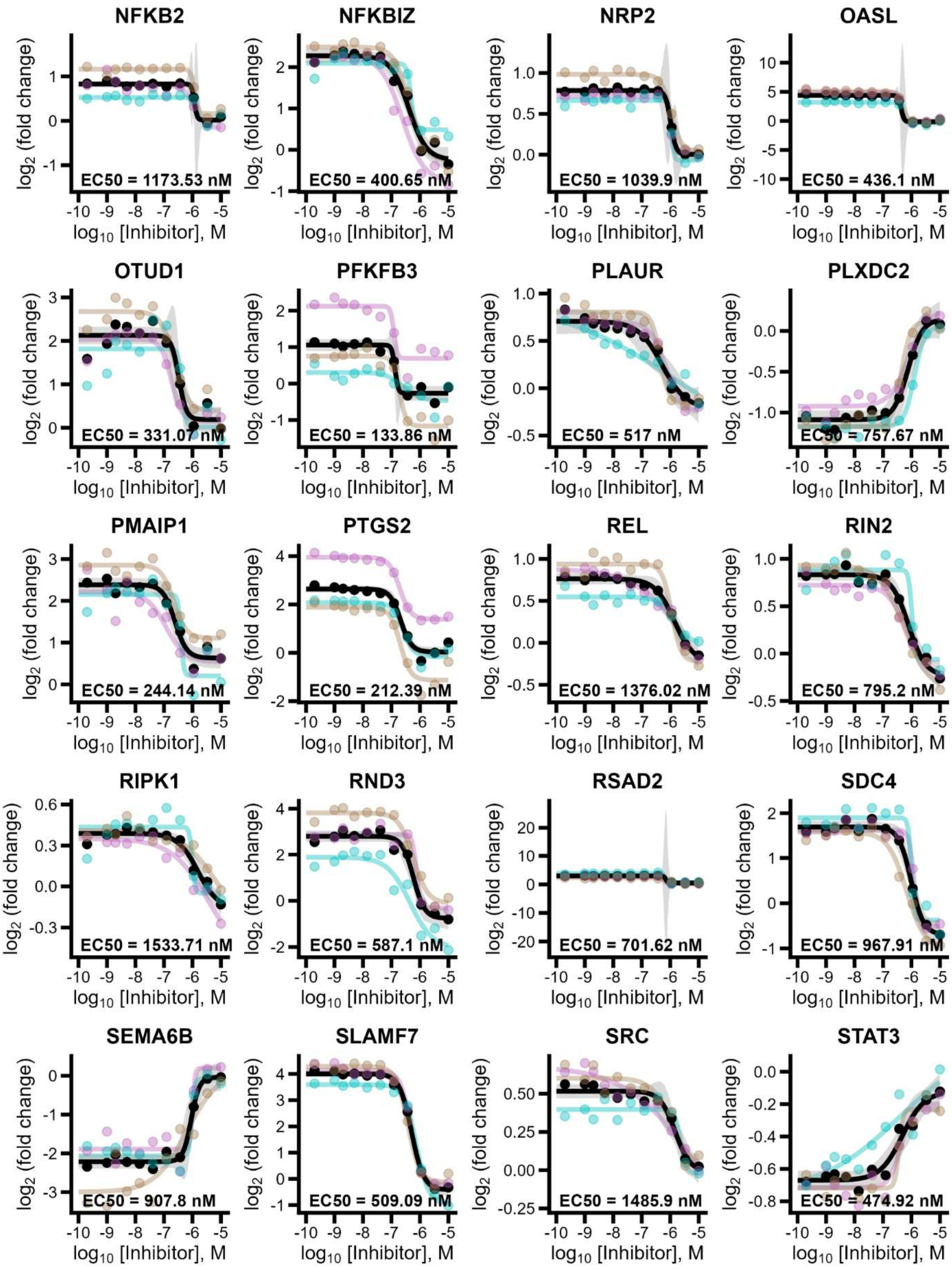
NFKB2
EC50 = 1173.53 nM
NFKBIZ
EC50 = 400.65 nM
NRP2
EC50 = 1039.9 nM
OASL
EC50 = 436.1 nM
OTUD1
EC50 = 331.07 nM
PFKFB3
EC50 = 133.86 nM
PLAUR
EC50 = 517 nM
PLXDC2
EC50 = 757.67 nM
PMAIP1
EC50 = 244.14 nM
PTGS2
EC50 = 212.39 nM
REL
EC50 = 1376.02 nM
RIN2
EC50 = 795.2 nM
RIPK1
EC50 = 1533.71 nM
RND3
EC50 = 587.1 nM
RSAD2
EC50 = 701.62 nM
SDC4
EC50 = 967.91 nM
SEMA6B
EC50 = 907.8 nM
SLAMF7
EC50 = 509.09 nM
SRC
EC50 = 1485.9 nM
STAT3
EC50 = 474.92 nM
$\log_2$ (fold change)
$\log_{10}$ [Inhibitor], M

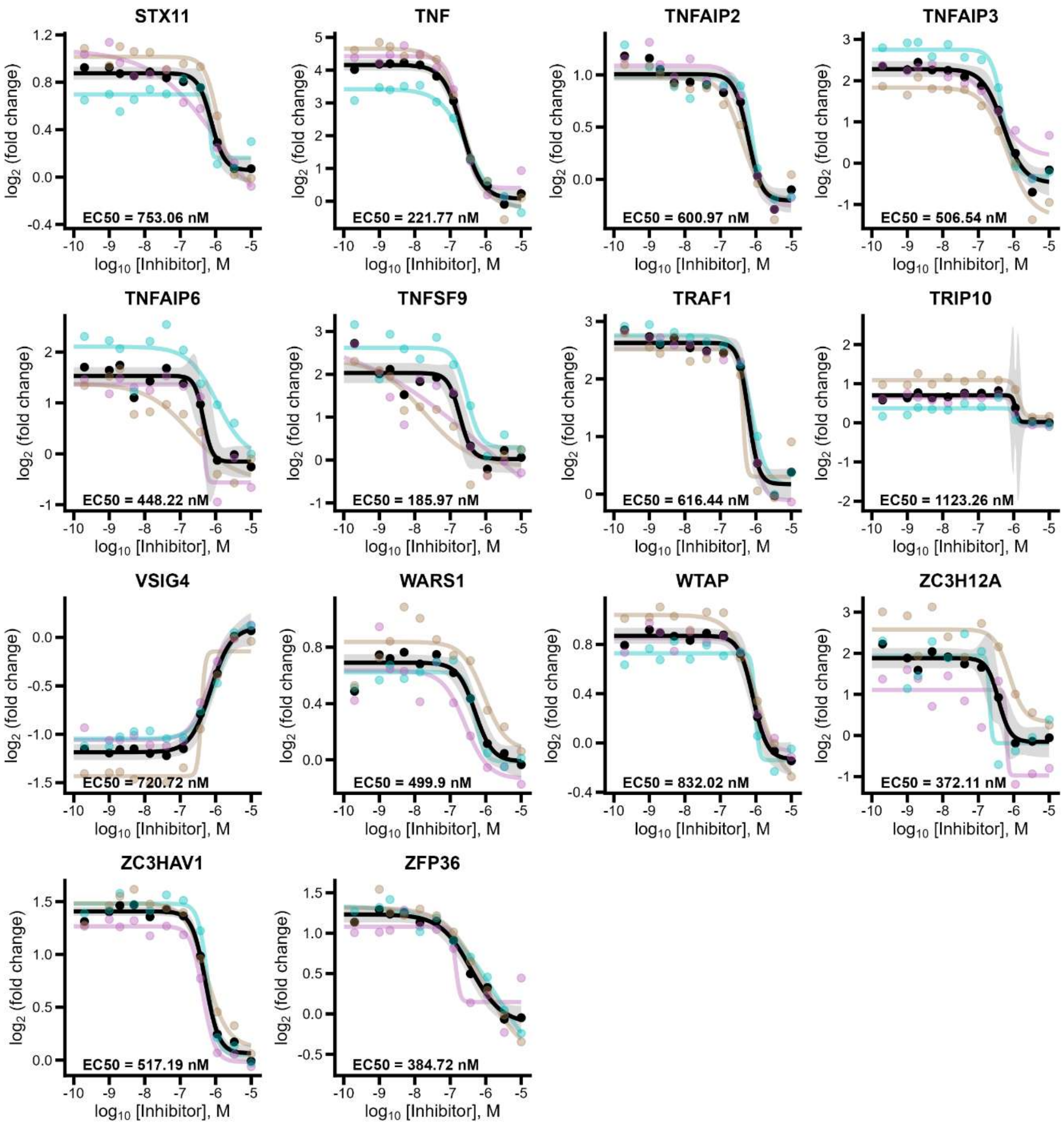


**Supplemental Fig. S8** Concentration-response curves showing $log_2$(fold change) in intracellular protein abundance relative to M0 controls across an 11-point inhibitor concentration range (0.16 nM – 10 µM) for TAK-242 (TLR4 inhibitor) Coloured curves represent fits to individual donors (n = 3), with corresponding data points shown, while the black curve represents the fit to combined data across donors (n = 9). Shaded regions indicate 95% confidence intervals. $EC_{50}$ values calculated from the combined fit are indicated for each protein. Selected proteins passed curve-fitting criteria and exhibit concentration-dependent changes in abundance following TLR4 inhibition.

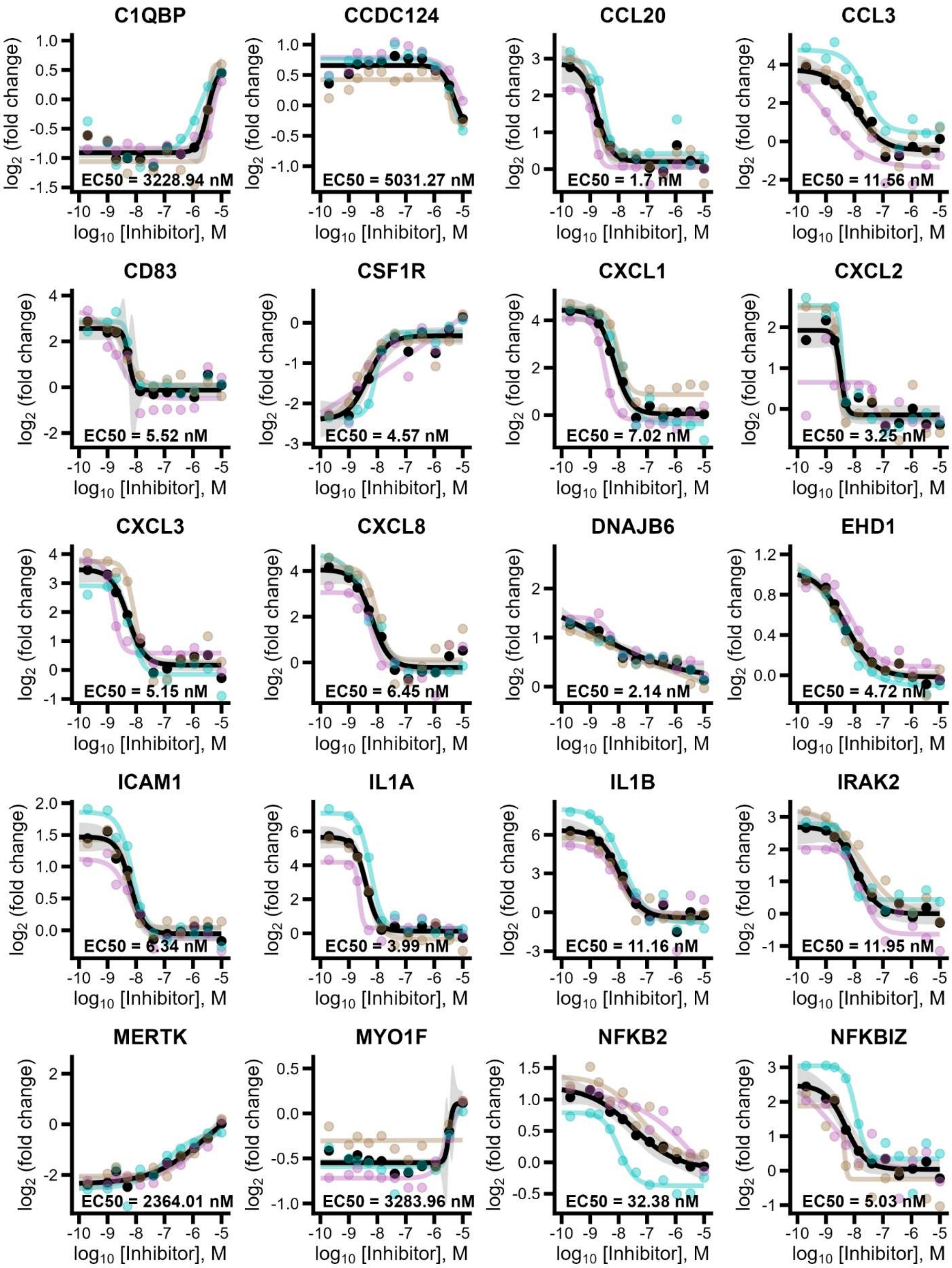
C1QBP
EC50 = 3228.94 nM
CCDC124
EC50 = 5031.27 nM
CCL20
EC50 = 1.7 nM
CCL3
EC50 = 11.56 nM
CD83
EC50 = 5.52 nM
CSF1R
EC50 = 4.57 nM
CXCL1
EC50 = 7.02 nM
CXCL2
EC50 = 3.25 nM
CXCL3
EC50 = 5.15 nM
CXCL8
EC50 = 6.45 nM
DNAJB6
EC50 = 2.14 nM
EHD1
EC50 = 4.72 nM
ICAM1
EC50 = 6.34 nM
IL1A
EC50 = 3.99 nM
IL1B
EC50 = 11.16 nM
IRAK2
EC50 = 11.95 nM
MERTK
EC50 = 2364.01 nM
MYO1F
EC50 = 3283.96 nM
NFKB2
EC50 = 32.38 nM
NFKBIZ
EC50 = 5.03 nM
log2 (fold change)
log10 [Inhibitor], M

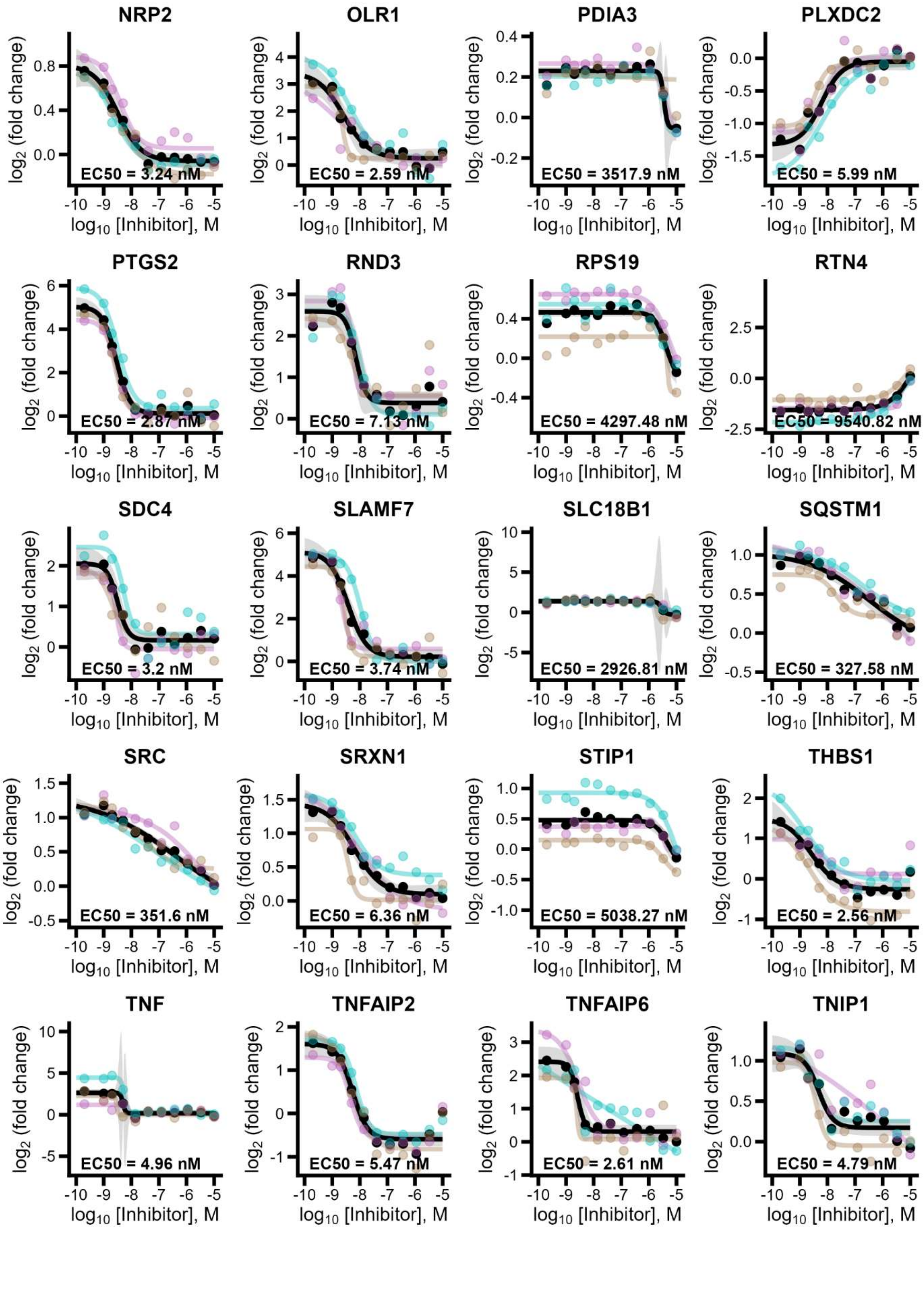
NRP2
EC50 = 3.24 nM
OLR1
EC50 = 2.59 nM
PDIA3
EC50 = 3517.9 nM
PLXDC2
EC50 = 5.99 nM
PTGS2
EC50 = 2.87 nM
RND3
EC50 = 7.13 nM
RPS19
EC50 = 4297.48 nM
RTN4
EC50 = 9540.82 nM
SDC4
EC50 = 3.2 nM
SLAMF7
EC50 = 3.74 nM
SLC18B1
EC50 = 2926.81 nM
SQSTM1
EC50 = 327.58 nM
SRC
EC50 = 351.6 nM
SRXN1
EC50 = 6.36 nM
STIP1
EC50 = 5038.27 nM
THBS1
EC50 = 2.56 nM
TNF
EC50 = 4.96 nM
TNFAIP2
EC50 = 5.47 nM
TNFAIP6
EC50 = 2.61 nM
TNIP1
EC50 = 4.79 nM
log2 (fold change)
log10 [Inhibitor], M

**Supplemental Fig. S9** Concentration-response curves showing $log_2$(fold change) in intracellular protein abundance relative to M0 controls across an 11-point inhibitor concentration range (0.16 nM – 10 µM) for MHV370 (TLR7/8 inhibitor). Coloured curves represent fits to individual donors (n = 3), with corresponding data points shown, while the black curve represents the fit to combined data across donors (n = 9). Shaded regions indicate 95% confidence intervals. $EC_{50}$ values calculated from the combined fit are indicated for each protein. Selected proteins passed curve-fitting criteria and exhibit concentration-dependent changes in abundance following TLR7/8 inhibition.
